# A quantitative mapping approach to identify direct interactions within complexomes

**DOI:** 10.1101/2021.08.25.457734

**Authors:** Philipp Trepte, Christopher Secker, Soon Gang Choi, Julien Olivet, Eduardo Silva Ramos, Patricia Cassonnet, Sabrina Golusik, Martina Zenkner, Stephanie Beetz, Marcel Sperling, Yang Wang, Tong Hao, Kerstin Spirohn, Jean-Claude Twizere, Michael A. Calderwood, David E. Hill, Yves Jacob, Marc Vidal, Erich E. Wanker

## Abstract

Complementary methods are required to fully characterize all protein complexes, or the complexome, of a cell. Affinity purification coupled to mass-spectrometry (AP-MS) can identify the composition of complexes at proteome-scale. However, information on direct contacts between subunits is often lacking. In contrast, solving the 3D structure of protein complexes can provide this information, but structural biology techniques are not yet scalable for systematic, proteome-wide efforts. Here, we optimally combine two orthogonal high-throughput binary interaction assays, LuTHy and N2H, and demonstrate that their quantitative readouts can be used to differentiate direct interactions from indirect associations within multiprotein complexes. We also show that LuTHy allows accurate distance measurements between proteins in live cells and apply these findings to study the impact of the polyglutamine expansion mutation on the structurally unresolved N-terminal domain of Huntingtin. Thus, we present a new framework based on quantitative interaction assays to complement structural biology and AP-MS techniques, which should help to provide first-approximation contact maps of multiprotein complexes at proteome-scale.

**Graphical Abstract:** 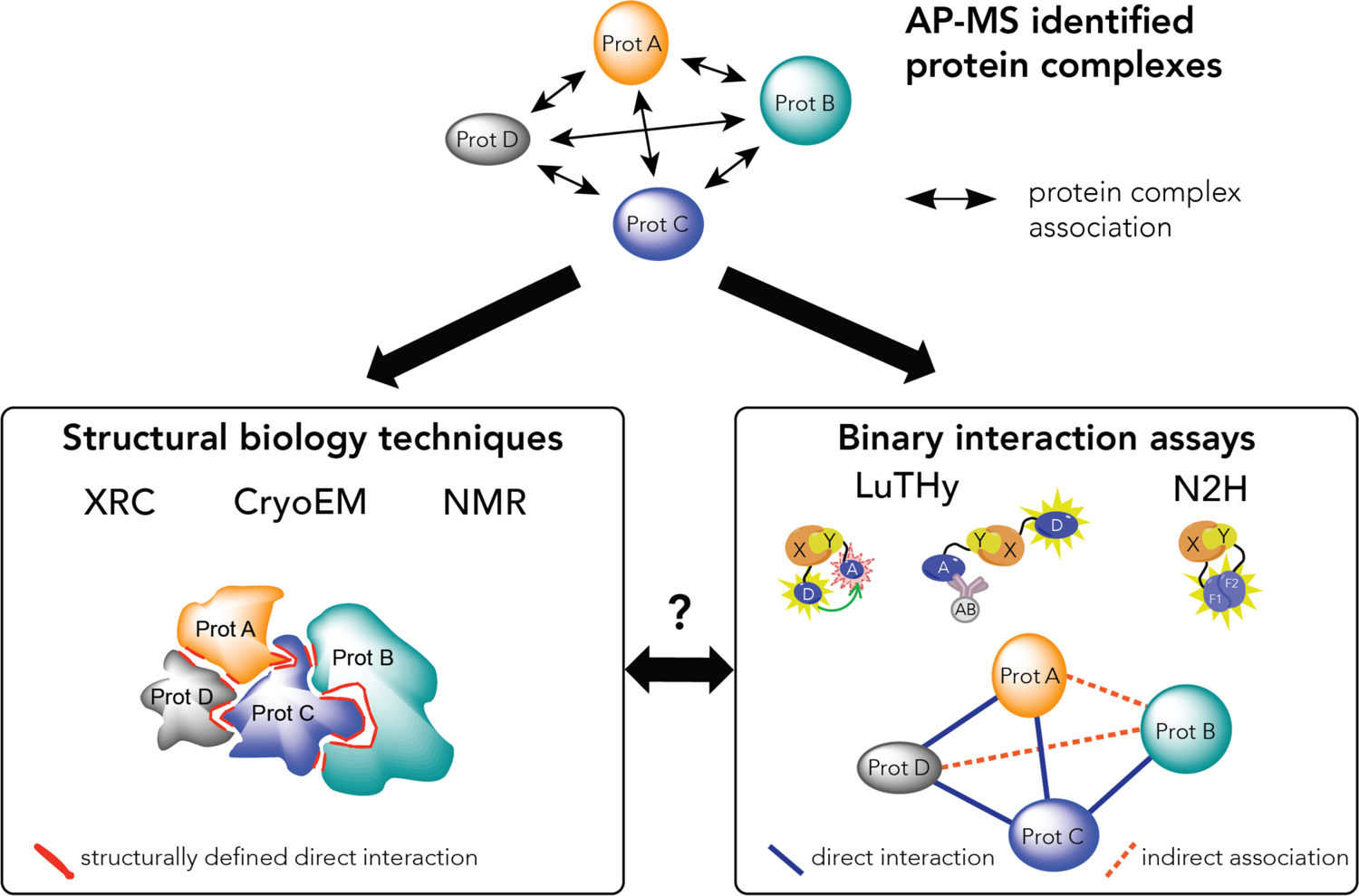

## INTRODUCTION

Characterizing the molecular architecture of all protein complexes, or the complexome, of a cell is essential to fully decipher its biological functions^1, 2^. Protein complexes vary in sizes and compositions as they often consist of different types of macromolecules including DNA, RNA and lipids. For example, in human cells, ∼80 proteins closely assemble with different ribosomal RNAs to form a functional ribosome, which is a compact spherical particle with a diameter of 250 to 300 Å^3, 4^. In comparison, the human proteasome is composed of eight protein subunits that co-assemble into a stable cylindrical complex^5^. This exemplifies how structurally diverse proteins can dynamically assemble in cells to perform various biological functions.

Affinity purification coupled to mass spectrometry (AP-MS) techniques are highly efficient in identifying the composition of protein complexes at proteome-scale^6–9^, but their ability to elucidate direct contacts between potentially interacting subunits is limited^10, 11^. However, gaining information on direct interactions, i.e. on proteins sharing a physical interface, and indirect associations, i.e. proteins that do not directly interact, within complexes is important to understand their biological roles. For example, clearly defining interacting domains between proteins will help in the design of hypothesis-driven functional studies. Additionally, defining the relationship between complex members is crucial to characterize the effect of disease-associated missense mutations on protein assemblies, or for the identification and validation of drug targets.

Structural biology technologies such as X-ray diffraction (XRD), nuclear magnetic resonance (NMR), or cryo-electron microscopy (cryo-EM) provide high-resolution data on the assembly of multiprotein complexes. This includes information on the folding of protein domains, amino acid composition at contact sites, attachment of compounds to proteins, and post-translational modifications of specific amino acids. However, in comparison to high-throughput AP-MS-based methods, generation of protein interaction data using structural biology techniques can be labor intensive and time consuming. In addition, such techniques often use truncated instead of full-length proteins, and comprehensive, high-resolution structural information is still missing for most protein complexes. Indeed, based on a recent study by Drew et al, ∼7,000 human protein complexes can be found in the human proteome from the integration of over 15,000 published mass-spectrometry experiments^12, 13^. However, only ∼4% of those (i.e. 309 protein complexes) currently have a resolved structure in the literature. Thus, complementary techniques are required to precisely characterize protein complexes currently identified by AP-MS-based methods^7, 11, 14^.

Previous proteome-scale investigations of binary protein-protein interactions (PPIs) revealed that the yeast two-hybrid (Y2H) system preferentially detects directly interacting proteins^10, 15^, and can thus provide a first-approximation view of the overall organization of complexes. While relatively versatile and amenable to high-throughput strategies, binary interaction assays are somewhat limited by their sensitivity^16, 17^. We have recently demonstrated that this limitation can be overcome by optimally combining complementary assays and/or versions thereof to increase sensitivity without affecting specificity^18, 19^.

Here we investigate a new combination of two binary interaction assays, the bioluminescence-based two-hybrid (LuTHy) method^19^, and the mammalian cell expression version of the NanoLuc two-hybrid (mN2H) assay^18^ to systematically map pairwise interactions between subunits of three multiprotein complexes, LAMTOR, BRISC and MIS12, for which both AP-MS^7, 11, 14^ and high-resolution structural data are available^20–22^. We find that direct interactions confer significantly higher scores than indirect associations, which allows their differentiation. Finally, we provide experimental evidence that the bioluminescence resonance energy transfer (BRET) readout of LuTHy can be used as a molecular ruler in live cells to estimate distances between protein domains for which no high-resolution structure is currently available. Our combined quantitative PPI mapping approach should thus be applicable for the in-depth characterization of subunit interactions at complexome-scale.

## RESULTS

### Benchmarking LuTHy under maximized specificity

We previously demonstrated that combining multiple complementary interaction assays and/or versions thereof significantly increases PPI recovery^17–19^. Here, we first benchmarked the PPI detection performance of LuTHy, a bioluminescence-based double-readout technology^19^, against an established positive reference set (PRS), hsPRS-v2, which contains 60 well-characterized human PPIs^17, 18^. To control for specificity, a random reference set (RRS), hsRRS-v2, made of 78 pairs of human proteins not known to interact, was also tested^17, 18^. With LuTHy, the interaction between two proteins of interest, X and Y, is first measured in living cells by quantification of the BRET signal (LuTHy-BRET; **Supplementary Figure 1A**). Then, following cell lysis, the same interaction is assessed *in vitro* by a quantitative luminescence-based co-precipitation readout (LuTHy-LuC; **Supplementary Figure 1A**). Since LuTHy plasmids allow expression of each protein as N- or C-terminal fusions, and as donor (NL; NanoLuc tag) or acceptor (mCit; mCitrine tag) proteins, eight tagging configurations can be assessed for every protein pair of interest (**Supplementary Figure 1B**). Thus, when all eight configurations are tested, a typical LuTHy experiment with both BRET and LuC measurements generates a total of 16 data points (**Figure 1A,B; Source Data Figures 1-2**).

**Figure 1.**
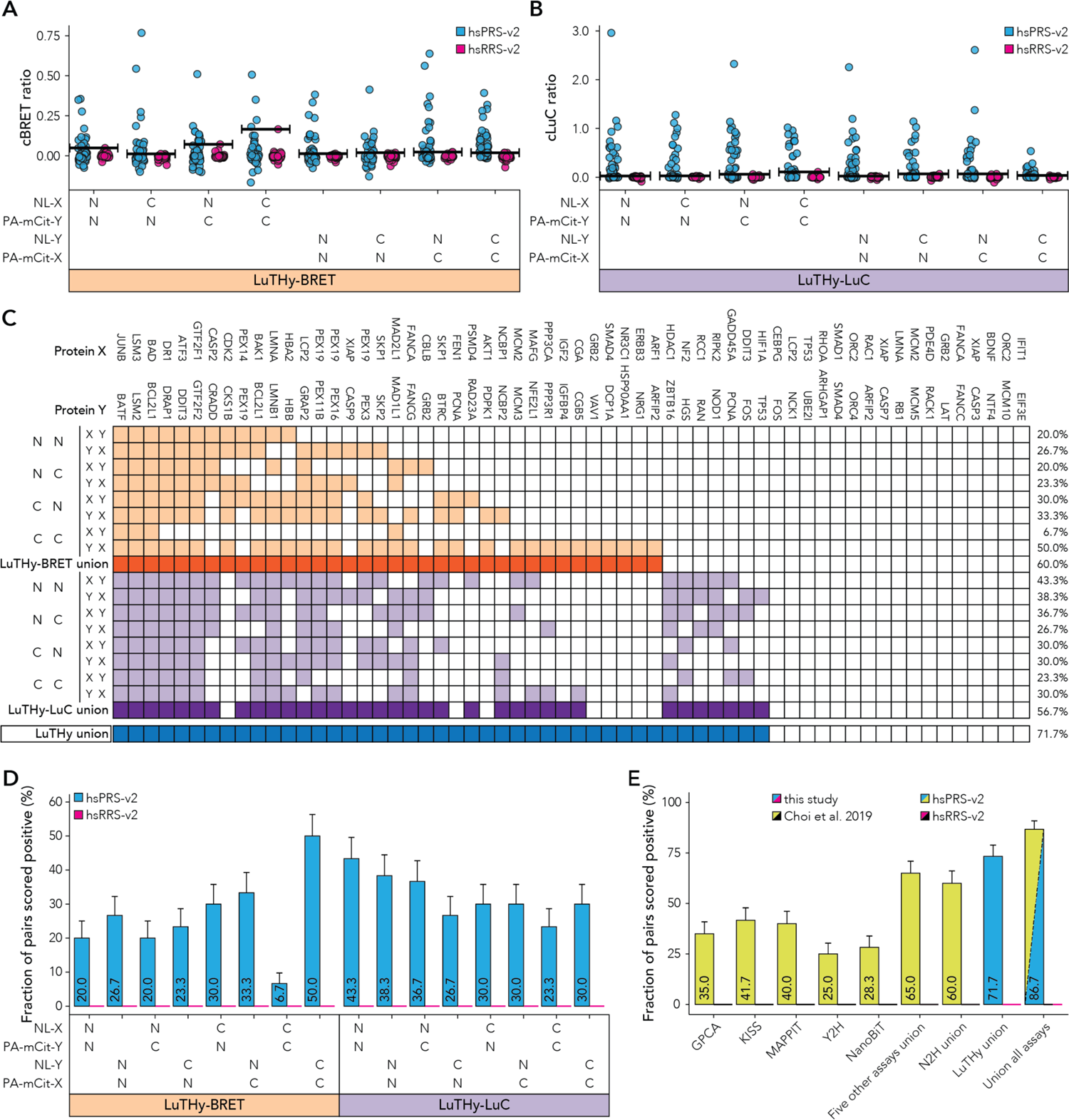
Benchmarking the LuTHy assay against hsPRS-v2 under conditions where none of the hsRRS-v2 pairs are scored positive. Quantitative scores for hsPRS-v2 and hsRRS-v2 pairs differentiated by tagging configurations for (**A**) LuTHy-BRET and (**B**) LuTHy-LuC. Horizontal lines in (**A**) and (**B**) indicate the scoring cutoffs for each configuration above which no hsRRS-v2 pair is scored positive. (**C**) Overview of the positively scored interactions from hsPRS-v2. Positive LuTHy-BRET interactions in different orientations (light orange) and their union (dark orange) are shown, as well as positive LuTHy-LuC interactions in the eight different configurations (light purple) and their union (dark purple). LuTHy union (blue) summarizes results for all 16 versions. The percentage (%) at the end of each row represents the fraction of hsPRS-v2 PPIs scored positive. (**D**) Recovery rates of hsPRS-v2 interactions at no hsRRS-v2 detection for the eight different tagging configurations in LuTHy-BRET and LuTHy-LuC. (**E**) Comparison of hsPRS-v2 recovery rates for LuTHy (LuTHy union) to other binary interaction methods benchmarked in Choi et al^18^. Error bars indicate standard errors of the proportion in (**D**) and (**E**).

**Figure 2.**
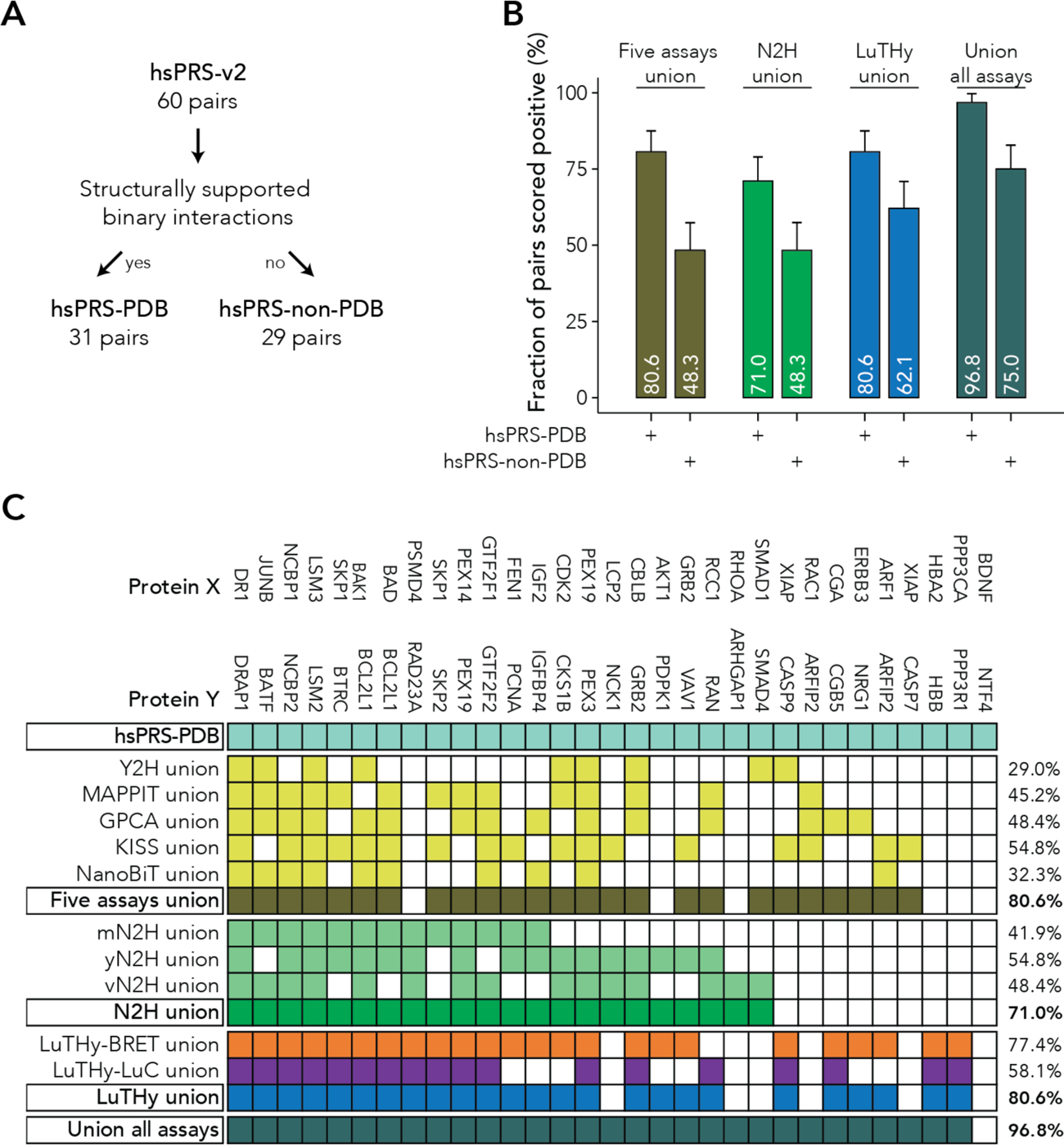
Benchmarking binary PPI assays against structurally supported interactions in hsPRS-v2. (**A**) hsPRS-v2 contains 60 binary PPIs among which 31 pairs are supported by structural data in PDB (hsPRS-PDB). The 29 remaining pairs are not currently supported by 3D structures (hsPRS-non-PDB). (**B**,**C**) Comparison of hsPRS-PDB and hsPRS-non-PDB recovery rates from different binary interaction methods when none of the hsPRS-v2 pairs are scored positive. Binary PPI assays combined (“Union all assays”) recover all but one of the hsPRS-PDB interactions. The percentage (%) at the end of each row in (**C**) indicates the fraction of hsPRS-v2 PPIs scored positive at no hsRRS-v2 detection. Error bars in (**B**) indicate standard errors of the proportion.

PPI detection in hsPRS-v2 by LuTHy was measured under conditions of maximal specificity, i.e. under conditions where none of the random protein pairs from hsRRS-v2 are scored positive (**Supplementary Figure 1C**). In total, using all eight tagging configurations with both readouts (i.e. BRET and LuC), LuTHy identified 43 of the 60 PPIs tested in hsPRS-v2, i.e. 71.7% (**Figure 1C-E**), confirming our previous observation that binary PPI mapping can be maximized when multiple assay versions are combined^18, 19^.

As reported in Choi et al^18^, under identical conditions of maximal specificity, the union of 16 versions of five distinct PPI assays, GPCA, KISS, MAPPIT, Y2H and NanoBiT, identified 65.0% of the hsPRS-v2 PPIs, while the union of 12 N2H assay versions reached a detection of 60.0% (**Figure 1E; Supplementary Figure 2A,B**). Thus, LuTHy, with its double readout system, is a highly sensitive and specific PPI detection method that recovers most of the well-established hsPRS-v2 interactions.

### Detecting direct interactions supported by high-resolution structural data

To further evaluate the extent to which different binary PPI assays can detect directly interacting proteins, we compared the recovery of the 31 hsPRS-v2 PPIs for which high-resolution structures are available in the RCSB protein data bank (PDB)^23^ (hsPRS-PDB), to that of the 29 remaining interactions (hsPRS-non-PDB) (**Figure 2A**). Our analysis revealed that structurally supported PPIs are detected with higher success rates in comparison to interactions for which structural information is not currently available (**Figure 2B**). This was the case for: 1) our previous benchmarking of five binary assays combined (Five assays union, 80.6% hsPRS-PDB vs. 48.3% hsPRS-non-PDB); 2) the union of 12 N2H assay versions (N2H union, 71.0% vs. 48.3%); and 3) the newly benchmarked LuTHy method (LuTHy union, 80.6% vs. 62.1%) (**Figure 2B**). Again, we confirmed our previous finding that the success of interaction detection significantly increases when multiple complementary PPI detection assays, or assay versions, are used. Indeed, we found that combining all seven PPI assays (GPCA, KISS, MAPPIT, Y2H, NanoBiT, N2H and LuTHy) allowed recovering 30 of the 31 structurally supported PPIs (i.e. 96.8%) under conditions of maximal specificity (Union all assays, **Figure 2B,C**). This demonstrates that structurally supported, direct PPIs can be recovered with nearly complete success rate when combining currently available binary PPI assays.

### Systematically mapping interactions within distinct multiprotein complexes

To generalize these findings, we decided to map PPIs between subunits of well-characterized human multiprotein complexes. We selected three complexes based on the following criteria: 1) identified by AP-MS-based methods^7, 11, 14^; 2) contains at least four subunits; 3) at least one 3D structure available in PDB^23^; and 4) at least 80% of cloned open reading frames (ORFs) encoding the reported subunits present in the human ORFeome 8.1 collection^24^. This resulted in a list of 24 distinct protein complexes (**Supplementary Table 1**), among which three structurally diverse candidates with well-characterized biological functions were prioritized: 1) the LAMTOR complex, also termed “Ragulator” complex^20^, which regulates MAP kinases and mTOR activities and consists of seven subunits (LAMTOR1, LAMTOR2, LAMTOR3, LAMTOR4, LAMTOR5, RRAGA and RRAGC); 2) the MIS12 complex that connects the kinetochore to microtubules^21^, and is made of five subunits (CENPC1, DSN1, MIS12, NSL1 and PMF1); and 3) the BRISC complex, a large deubiquitinating machinery^22^ consisting of five proteins (ABRAXAS2, BABAM1, BABAM2, BRCC3 and SHMT2) (**Figure 3A, Supplementary Table 3**).

**Figure 3.**
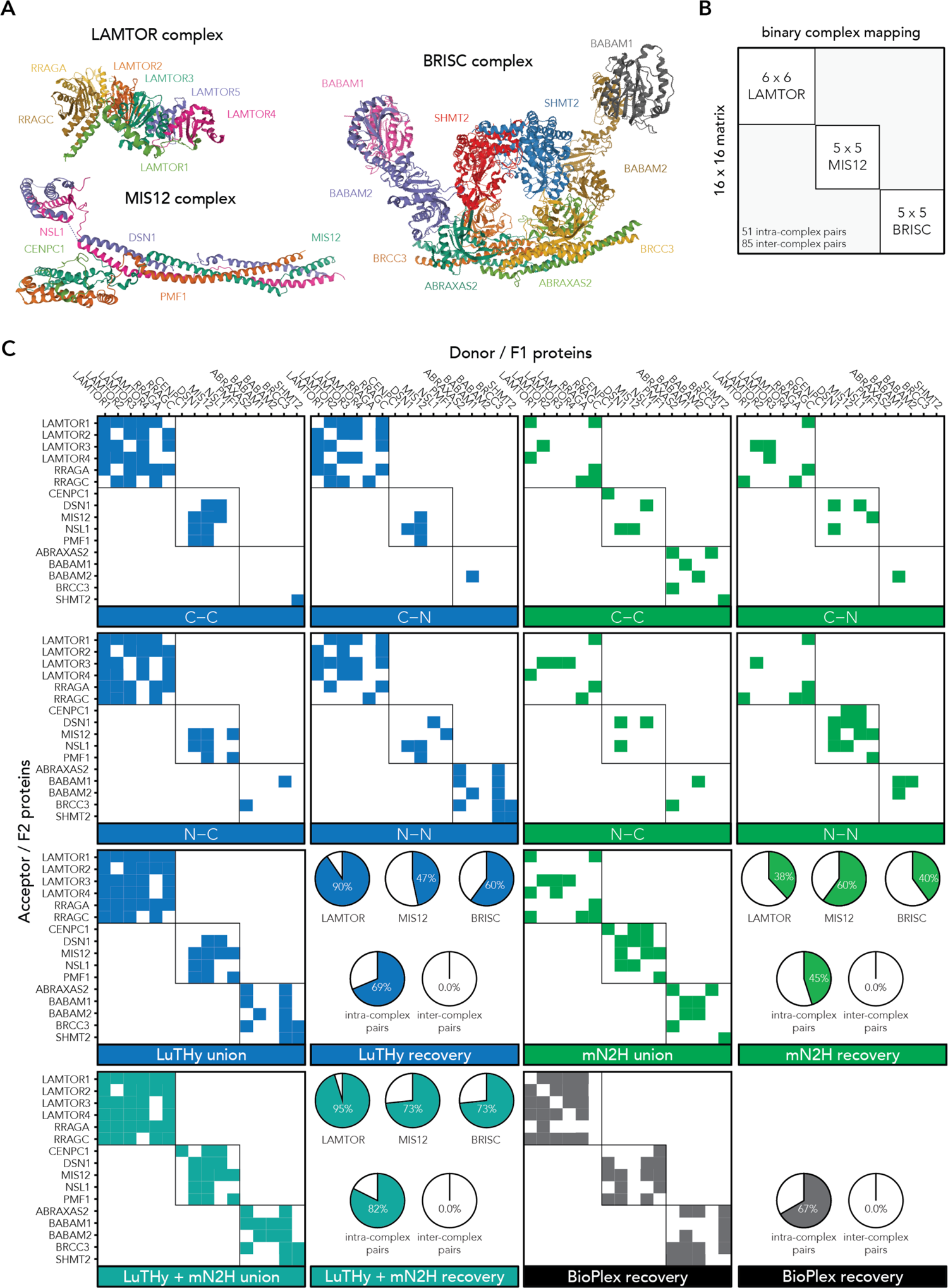
Mapping interactions within multiprotein complexes using the LuTHy and mN2H assays. (**A**) Structures of the protein complexes analyzed in this study: LAMTOR (PDB: 6EHR), MIS12 (PDB: 5LSK), and BRISC (PDB: 6H3C). (**B**) Binary interaction approach to systematically map PPIs within distinct complexes. Every protein subunit from each complex was screened against every other one (all-by-all, 16×16 matrix). (**C**) Results of the all-by-all interaction screen for the selected multiprotein complexes. Each protein pair was systematically tested in every possible configuration (i.e. C-C, C-N, N-C, N-N), and as donor and acceptor in LuTHy, or as F1 and F2 NanoLuc fusions in mN2H. LuTHy union corresponds to the combination of both LuTHy-BRET and LuTHy-LuC results. Protein pairs tested in the LuTHy assays were scored positive if at least one of the two readouts (i.e. LuTHy-BRET and/or LuTHy-LuC) was positive. LuTHy + mN2H union corresponds to the combination of results from LuTHy union and mN2H union. In the pie charts, the top panels show recovery rates of intra-complex pairs within the LAMTOR, MIS12 and BRISC complexes, while the bottom panels show recoveries of all intra-complex and inter-complex pairs. Published BioPlex results, corresponding to AP-MS data, were used for the interaction matrix and pie charts. Proteins used as baits in BioPlex are indicated on the x-axis, whereas identified interaction partners are indicated on the y-axis.

To map interactions between subunits of the LAMTOR, MIS12 and BRISC complexes, we systematically tested all possible pairwise combinations with the LuTHy and mN2H assays (**Source Data Figures 3-5**). Except for LAMTOR5, which was not available in the human ORFeome 8.1 collection, all 16 ORFs encoding the selected target proteins were sequence verified and cloned into both LuTHy and N2H expression plasmids. A resulting search space of 256 pairwise combinations, corresponding to a total of 16 subunits for the three complexes (16×16 matrix; **Figure 3B**), was thus systematically explored with the LuTHy and mN2H assays. To score a tested interaction as positive, we rationalized that true binary PPIs should only be found between the respective subunits of a given complex (i.e. intra-complex pairs), but not between subunits belonging to different complexes (i.e. inter-complex pairs). Therefore, we treated all inter-complex pairs as negative controls, similar to protein pairs from a random reference set. We observed that LuTHy and mN2H fusion constructs showed a broad distribution of interaction scores among those inter-complex pairs (**Supplementary Figure 3A-F**). Therefore, we defined construct-specific cutoffs under conditions of maximal specificity, where none of the inter-complex protein pairs are scored positive with any of the assay versions. Using this strategy, we found that LuTHy and mN2H recovered 35 (68.6%) and 23 (45.1%) of the 51 intra-complex interactions, respectively, while 42 of those protein pairs (82.4%) were detected when combining both assays (**Figure 3C; Supplementary Figure 4**). We then compared those results to the ones from the BioPlex dataset^7, 11, 14^. According to BioPlex, 66.7% of all intra-complex interactions but none of the inter-complex protein pairs for the LAMTOR, BRISC and MIS12 complexes were detected by AP-MS (**Figure 3C**). This demonstrates that, similar to AP-MS-based techniques, binary PPI mapping with LuTHy and mN2H can recover a large fraction of interactions between subunits of distinct multiprotein complexes with high sensitivity and specificity.

### Differentiating direct interactions from indirect associations

Differentiating direct interactions from indirect associations within each multiprotein complex identified by AP-MS-based methods is an important, unresolved challenge. In fact, it was estimated that only ∼32% of the detected protein associations in the BioPlex dataset are direct interactions^11^. Therefore, we evaluated if the binary PPI detection assays, LuTHy and mN2H, could identify directly interacting subunits within multiprotein complexes. To define direct interactions and indirect associations between the subunits of each complex, we used 3D structural information available in three public databases: PDB^23^, Interactome3D^25^ and PDBePISA^26^. While directly interacting proteins share a physical interface, we defined indirect associations as pairs of subunits that do not share an interaction interface (**Figure 4A**). In total, we classified 31 direct interactions and 7 indirect associations between the subunits of the LAMTOR, BRISC and MIS12 complexes. Noticeably, we excluded 13 separate homodimer interactions from all further analyses since they were not reported in the available multiprotein complex structures (**Supplementary Table 2**). When applying the same cutoffs used to differentiate intra-from inter-complex protein pairs, 70.9%, 45.2%, and 48.4% of the direct interactions could be detected with LuTHy-BRET, LuTHy-LuC, and mN2H, respectively (**Supplementary Figure 5A)**. While mN2H only detected direct interactions, LuTHy-BRET and LuTHy-LuC also recovered 71.4% and 28.6% of the indirect associations, respectively (**Supplementary Figure 5B**). The union of LuTHy and mN2H detected a total of 80.6% of the structure-based direct interactions, and 85.7% of the indirect associations (**Supplementary Figure 5A,B**). In comparison, according to the BioPlex dataset, 87.1% of the direct interactions along with 100% of the indirect associations were recovered by AP-MS (**Supplementary Figure 5A,B**).

**Figure 4.**
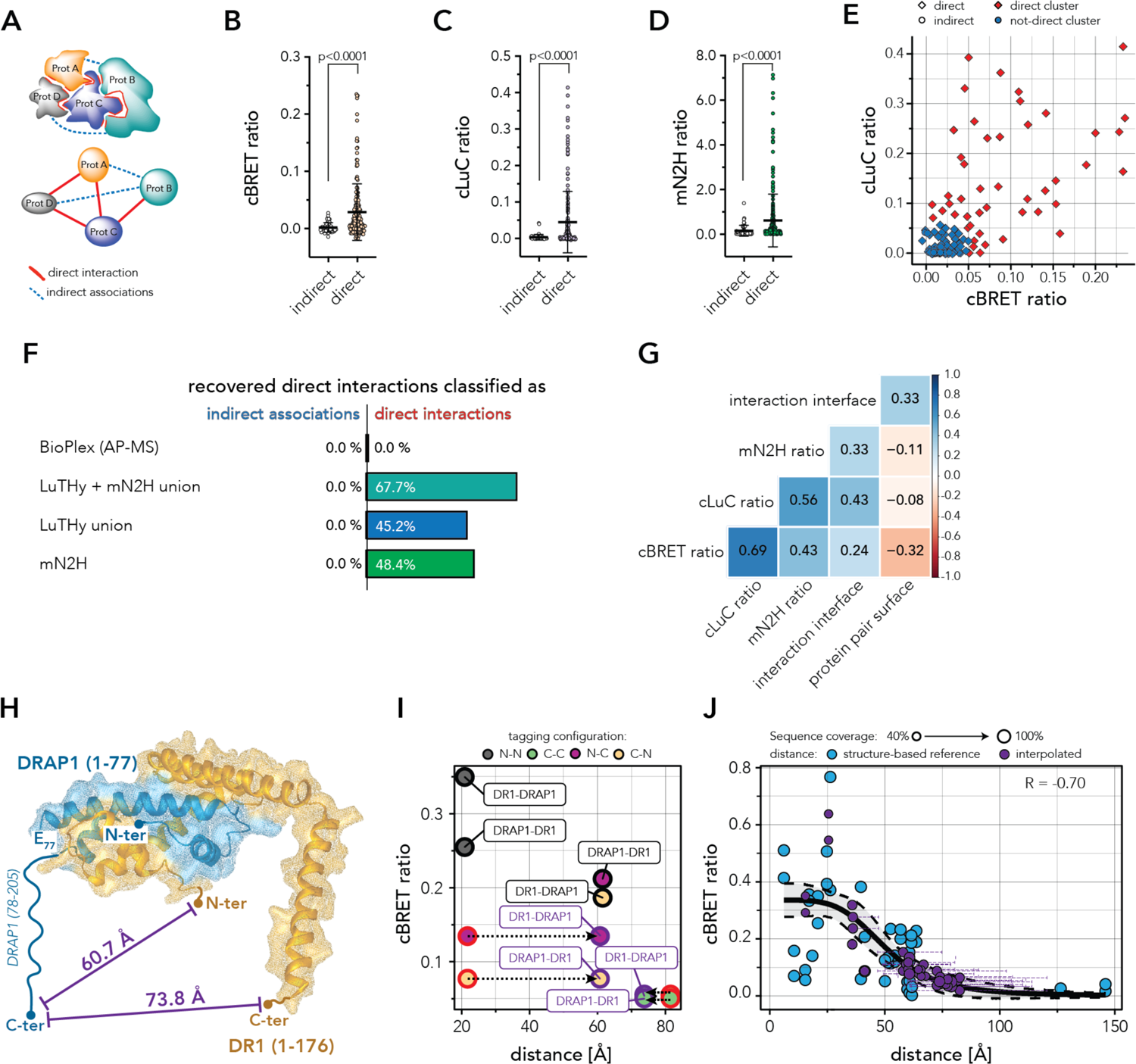
Differentiating direct interactions from indirect associations with LuTHy and mN2H assays. (**A**) Schematic for the definition of direct interactions and indirect associations within complexes using structural biology data. (**B-D**) Distribution of the quantitative scores and comparison between direct interactions and indirect associations. Direct interactions produce significantly higher LuTHy-BRET (**B**), LuTHy-LuC (**C**), and mN2H (**D**) scores compared to indirect associations (Welch’s two-tailed t-test). (**E**) Scatter plot of cBRET and cLuC ratios showing direct interactions (diamonds) and indirect associations (circles) detected by LuTHy-BRET or LuTHy-LuC. Protein pairs were clustered by supervised expectation-maximization clustering as direct (red) or not-direct (blue) interactions using hsPRS-PDB and hsRRS-v2 as training sets. (**F**) Recovery and classification of structurally defined direct interactions as true direct interactions or as indirect associations by BioPlex (AP-MS), LuTHy and mN2H. (**G**) Pearson correlation matrix for direct interactions comparing cBRET, cLuC and mN2H ratios to the interaction interface areas (Å^2^), or to the total complex surface areas (Å^2^). (**H**) 3D structure of the DR1-DRAP1 interaction (PDB: 1JFI). Full-length DR1 but only the N-terminal region of DRAP1 (1-77, amino acids 78-205 lacking) are reported in the 3D structure. The cBRET ratios were used to interpolate distances between the C-terminus of DRAP1 and the N-terminus (60.7 Å), or C-terminus (73.8 Å) of DR1 based on the cBRET-distance standard curve. (**I**) cBRET ratios for the DR1-DRAP1 PPI are plotted against the molecular distances obtained from the 3D structure. Tagging configurations are colored by N-N (grey), C-C (green), N-C (purple) or C-N (yellow). Each data point on the graph is labeled (framed text) according to the tested tagging configuration: the protein indicated first is tagged with NanoLuc (NL) luciferase, while the second protein is tagged with PA-mCitrine (PA-mCit) (e.g. DR1+DRAP1 (N-N, grey) corresponds to NL-DR1/PA-mCit-DRAP1). Tagging configurations where tags are fused to the structurally unresolved termini in the current 3D structure for one of the two proteins are outlined in red (e.g. DR1+DRAP1 (N-C, purple) corresponds to NL-DR1/DRAP1-mCit-PA). Interaction values outlined in purple represent cBRET ratios against interpolated molecular distances. Differences between structurally determined and interpolated distances are indicated by dotted arrows. (**J**) cBRET ratios plotted against the structurally determined molecular distances for the 44 protein pairs used as references (blue). The 30 PPIs with tags fused to protein termini not currently resolved in the structures are plotted against interpolated distances between full-length proteins (purple). For each tested pair, the purple horizontal error bar corresponds to the 95% confidence interval of the interpolated molecular distances. Sigmoidal fits were performed on the 44 tested pairs outlined in black, together with the 30 pairs outlined in purple (R = −0.70).

To further evaluate if we could confidently differentiate between direct interactions and indirect associations, we analyzed the quantitative scores obtained with LuTHy and mN2H. We observed that direct interactions generate significantly higher scores compared to indirect associations in the LuTHy-BRET (**Figure 4B**), LuTHy-LuC (**Figure 4C**) and mN2H (**Figure 4D**) assays. Since LuTHy recovered both direct interactions and indirect associations, we applied a machine learning-based clustering algorithm on the LuTHy-BRET and LuTHy-LuC scores to classify LuTHy-positive protein pairs as direct or not-direct interactions. We used the detected hsPRS-PDB PPIs as a positive training set for direct interactions, and the hsRRS-v2 pairs as a negative training set for not-direct interactions (**Supplementary Table 4**). Next, we applied the trained cluster algorithm to the 23 direct interactions and six indirect associations detected by LuTHy (**Supplementary Figure 5A,B)**. Using this unbiased approach, we were able to classify 14 of the LuTHy-positive direct interactions (45.2%) as true direct PPIs, without wrongly classifying any of the indirect associations (**Figure 4E,F, Supplementary Figure 5C**). After combining these results with the ones from the mN2H assay, we were able to confidently recover and classify 67.7% of the direct interactions among the LAMTOR, BRISC and MIS12 complexes. In comparison, protein pair associations reported in the BioPlex database are currently neither classified as direct PPIs or indirect associations (**Figure 4F, Supplementary Figure 5C**)^7, 11, 14^.

To better understand why a broad range of quantitative scores is obtained when the same pair of directly interacting proteins is tested in eight different tagging configurations (**Supplementary Figure 6A-I**), we explored the structural features of all direct interactions among the three complexes. Using PDBePISA^26^ we obtained the sizes of interaction interfaces as well as the total surface areas for all directly interacting subunits within the LAMTOR, BRISC and MIS12 complexes (**Supplementary Table 5**). We observed a stronger correlation between interaction scores and the sizes of the interaction interfaces, rather than to the total surface areas of the protein pairs (**Figure 4G; Supplementary Figure 7A-F**), indicating that the larger the interaction interface, i.e. the stronger the binding^27^, the higher the interaction score. Noticeably, LuTHy-BRET showed relatively high corrected BRET (cBRET) ratios for small interaction interfaces (<500 Å^2^, **Supplementary Figure 7A**), which supports our previous findings that low affinity interactions can be detected with this assay^19^. Surprisingly, a significant negative correlation between cBRET ratios and the total surface areas of protein pairs was observed (**Figure 4G; Supplementary Figure 7D-F**), suggesting that the larger the co-complex, the lower the resonance energy transfer between the N- or C-terminally fused NanoLuc luciferase and mCitrine tags. This indicates that both affinity and distance between the tagged proteins studied with binary PPI assays can be determinants for the intensity of the quantitative scores, and thus for their detection.

To test whether the LuTHy-BRET readouts could be used to measure distances between protein domains, we determined the molecular distances between all protein termini among the three complexes using available structural information (**Source Data Figures 3-5**). We then compared those reference values to the measured cBRET ratios and found a significant negative correlation between them and the molecular distances of the protein termini (**Supplementary Figure 7G**). Interestingly, no such relationship could be observed for the non-energy transfer-based corrected LuC (cLuC) or mN2H ratios (**Supplementary Figure 7H,I**). In addition, the distances between protein termini of indirect associations identified with LuTHy-BRET were found to be within a similar range of distances than that of direct interactions (**Supplementary Figure 7J**). This suggests that LuTHy-BRET can detect indirect associations between proteins even in the absence of a common interaction interface as long as the tagged proteins are within a distance where an energy transfer between the donor and the acceptor can occur.

Altogether, these results show that the quantitative scores obtained with LuTHy and mN2H can be used to systematically identify direct interactions within multiprotein complexes and differentiate them from indirect associations. They also highlight that the measured cBRET ratios are correlated to the molecular distances between the protein termini of directly interacting subunits, which implies that LuTHy-BRET measurements could also be used to measure distances directly in live cells.

### Measuring molecular distances between subunits of protein complexes in live cells

Förster resonance energy transfer-based techniques have been previously used as spectroscopic rulers^28, 29^ to monitor conformational changes in proteins, or to estimate the proximity relationships of macromolecules in complexes^30–32^. Since high-resolution structural data are currently unavailable or incomplete for many protein complexes^13^, we investigated whether the LuTHy-BRET readout could be used to measure distances between full-length subunits of protein complexes in live cells. To establish a cBRET-distance standard curve, we used the structure-based molecular distances, the cBRET ratios of directly interacting subunits in the LAMTOR, BRISC and MIS12 complexes, and the structurally supported hsPRS-v2 PPIs (i.e. hsPRS-PDB) as references (**Supplementary Figure 8A, Source Data Figures 1-2** and **3-5**). However, many of these protein pairs are, on average, structurally covered by less than 75% of their respective full-length sequences (**Supplementary Figures 8B** and **9; Supplementary Tables 5** and **6, Source Data Figures 3-5**). Therefore, for the calibration, we selected a subset of 44 protein pairs (i.e. eight different interactions in multiple tagging-configurations) where the tagged protein termini are fully resolved in the associated structures. For this subset of structurally resolved protein pairs, we generated a standard curve by plotting the cBRET ratios against the measured distances. We observed a strong negative correlation between the cBRET ratios and the reported distances (R = −0.58), which is best described by a sigmoidal fit (**Supplementary Figure 8C**). This is in good agreement with previous reports describing that the energy transfer efficiency is inversely proportional to the sixth power of the distance between the donor and the acceptor^29^. This result strongly suggests that cBRET ratios can indeed be used to measure distances between protein domains in live cells.

We next compared cBRET ratios measured between full-length proteins *in cellulo*, to the reported distances in the corresponding high-resolution structures where truncated proteins were used. For example, in the structure of the DR1-DRAP1 PPI, all 176 amino acids of DR1 are resolved, while only the first 77 of the 205 amino acids of DRAP1 are structurally defined^33^. Thus, ∼62% of the DRAP1 protein, including its entire C-terminal domain, does not appear in the co-complex structure. The molecular distances reported in the structure suggest that the resolved termini of DRAP1 (N-terminus and glutamic acid 77) are both ∼20 Å apart from the N-terminus of DR1 (**Supplementary Figure 8D**). When comparing these reported distances to the LuTHy-BRET results, we observed higher cBRET ratios when DRAP1 and DR1 were both N-terminally tagged, compared to C-terminally fused DRAP1 and N-terminally fused DR1 (**Supplementary Figure 8E**). This suggests that the physical distance between the C-terminus of full-length DRAP1 and the N-terminus of DR1 is indeed larger in live cells than the ∼20 Å reported in the structure where a truncated DRAP1 protein was used. To estimate the distance between the structurally unresolved termini in the full-length co-complex, we inferred the distance from the LuTHy-BRET data by using the calibrated cBRET-distance standard curve (**Supplementary Figure 8C**). The measured cBRET ratios indicated that the DRAP1 C-terminus is ∼61 Å apart from the N-terminus, and ∼74 Å apart from the C-terminus of DR1 (**Figure 4H,I**). To generalize these findings, we extended the analysis to 30 additional PPIs that scored positive in LuTHy-BRET, and for which structural information on the tagged protein termini is currently missing (**Figure 4J**). As expected, we found that distances between full-length proteins in live cells inferred from LuTHy-BRET ratios do not generally match with the structurally reported distances (**Supplementary Figure 8F**, **Supplementary Figure 10**), confirming the results obtained for the DR1-DRAP1 interaction.

Together, these findings confirm that LuTHy-BRET measurements can provide valuable, complementary information about molecular distances between the subunits of protein complexes in live cells.

### Assessing disease mutation-induced dynamics in structurally unresolved protein domains of the Huntingtin-HAP40 interaction

To evaluate if the LuTHy-BRET readout could be used to analyze the effect of a disease mutation on the distance between protein domains, we studied the Huntington’s disease related Huntingtin-HAP40 complex. Huntington’s disease is caused by a CAG-repeat expansion located in exon-1 of the Huntingtin gene (*HTT*) that is translated into an elongated polyglutamine (polyQ) stretch within the first 83 amino acids of the >3000 amino acids HTT protein^34, 35^. Recently, the structure of full-length HTT, in complex with its binding partner HAP40, was solved at high-resolution by cryo-EM (**Figure 5A**)^36^. However, the polyQ stretch located at the N-terminus of HTT was not structurally solved, probably due to its high flexibility^36^. Thus, the structure and relative localization of the N-terminal, polyQ-containing domain remains elusive.

To investigate if the expansion mutation could influence the relative arrangement and molecular distance between the N-terminal, polyQ-containing domain of HTT and HAP40, we assessed this interaction by LuTHy-BRET. We observed a significant reduction in cBRET ratios when comparing the HTT-HAP40 interaction where HTT contains either a pathogenic (Q145), or a physiological (Q23) polyQ tract (**Figure 5B**). To determine the distance between the two proteins, we used the cBRET-distance standard curve (**Supplementary Figure 8C**) and found that the N-terminus of HAP40 is ∼53 Å apart from the N-terminus of HTT with the non-pathogenic Q23 tract (**Figure 5A; Supplementary Table 7**). However, the distance increases to ∼58 Å when HTT contains the elongated, pathogenic Q145 stretch (**Figure 5A; Supplementary Table 7**). This suggests that the polyQ expansion mutation alters the domain arrangements within the HTT-HAP40 complex, and that LuTHy-BRET is a powerful tool to measure such subtle changes in live cells.

**Figure 5.**
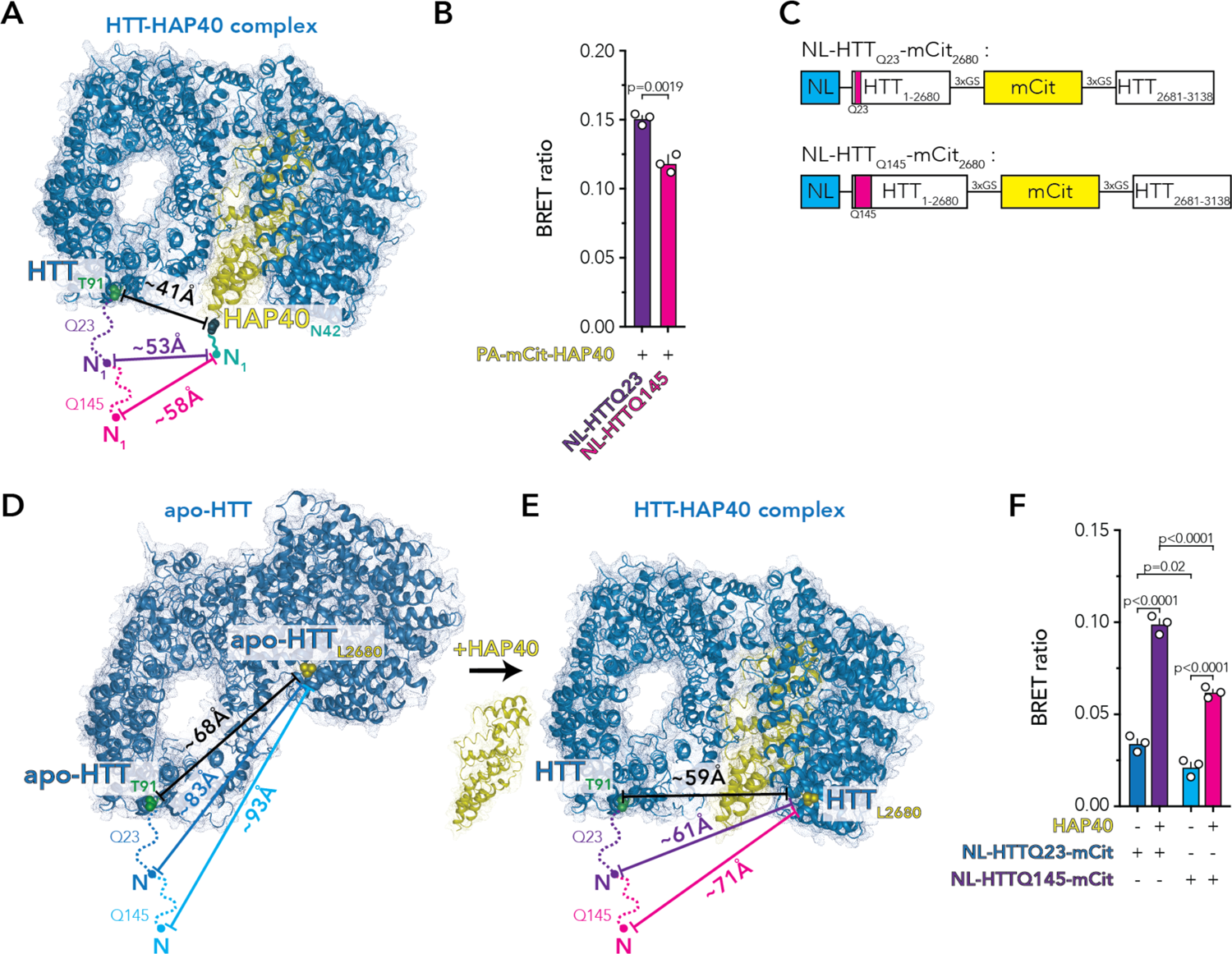
Interpolating molecular distances within the HTT-HAP40 complex with LuTHy-BRET. (**A**) Cryo-EM structure of the HTT-HAP40 complex (PDB: 6EZ8). The distances between HTT(T91) and HAP40(N42) inferred from the structure are indicated (black), along with the distances between the N-terminal domains of HAP40 and HTTQ23 (purple) or HTTQ145 (magenta) interpolated from the measured BRET ratios. (**B**) The BRET ratio between HAP40 and HTTQ23 is significantly higher than the one between HAP40 and HTTQ145 (the bar plots show means ± standard deviations (sd), statistical significance was calculated by a two-tailed t-test, n=3). (**C**) Schematic representations of the HTTQ23 and HTTQ145 intramolecular LuTHy-BRET sensors. (**D**,**E**) Cryo-EM structures of the apo-HTT protein (**D**, PBD: 6RMH) and the HTT-HAP40 complex (**E**, PDB: 6EZ8) where the distances between residues T91 and L2680 are indicated (black) either in the absence or presence of HAP40. (**F**) BRET ratios measured for the intramolecular LuTHy-BRET sensors in the absence or presence of HAP40, for HTTQ23 and HTTQ145 (the bar plots show means ± sd, statistical significance was calculated by two-way ANOVA with Tukey’s multiple comparisons test, n=3). The BRET ratios were used to interpolate distances between the N-terminus of HTT and residue L2680 for HTTQ23 and HTTQ145, which are indicated in **D** and **E**.

To assess if the effect of the elongated polyQ stretch on the relative arrangement of the N-terminal HTT domain is also detectable in the absence of HAP40, we analyzed the HTT protein in its apo form. The structure of the apo-HTT protein was recently solved and it was proposed that its C-terminal HEAT (Huntingtin/Elongation factor 3/protein phosphatase 2A/TOR) domain alters its position relative to the N-terminal HEAT domain upon polyQ expansion^37^. To monitor such a dynamic structural change within the HTT protein in live cells, we generated an intramolecular LuTHy-BRET sensor, where the NanoLuc (NL) luciferase is fused to the N-terminus, and the fluorescent acceptor, mCitrine (mCit), is incorporated into an unstructured loop at leucine 2680 (L2680, **Figure 5C-E**). We then quantified LuTHy-BRET signals for the resulting NL-HTTQ23-mCit(L2680) and NL-HTTQ145-mCit(L2680) sensor proteins (**Figure 5F**). Using the cBRET-distance standard curve, we interpolated that the HTTQ23 N-terminus is ∼83 Å apart from the loop at L2680, while this distance increases to ∼93 Å in the HTTQ145 protein (**Figure 5D; Supplementary Table 7**).

Interestingly, it was also shown that the apo-HTT conformation significantly differs from the HTT-HAP40 complex^37^. Upon HAP40 binding, the C-terminal HEAT domain undergoes a rearrangement, swinging into closer proximity to the N-terminus of HTT (**Figure 5E**). To monitor this dynamic rearrangement in live cells, we used the NL-HTTQ23-mCit(L2680) and NL-HTTQ145-mCit(L2680) LuTHy-BRET sensors and quantified the BRET ratios in presence or absence of exogenous HAP40 protein. When co-expressing HAP40 and the HTT sensors, the BRET ratio significantly increased, indicating that HTT also undergoes this structural rearrangement upon HAP40 binding in live cells (**Figure 5F**). In addition, in the presence of HAP40, the NL-HTTQ23-mCit(L2680) sensor showed a significantly higher BRET signal compared to the NL-HTTQ145-mCit(L2680) sensor, confirming that the polyQ expansion alters the domain arrangements of the HTT-HAP40 complex (**Figure 5E**). Using these data, we interpolated that in the presence of HAP40, the HTTQ23 N-terminus and the structural loop at L2680 are ∼61 Å apart, whereas they are ∼71 Å apart with the expanded polyQ tract (**Figure 5D; Supplementary Table 7**). Taken together, these results show that both in the presence or absence of HAP40, the pathogenic polyQ stretch-containing N-terminal domain of HTT is extending ∼10 Å further away from the core protein, a phenomenon that could not be resolved in the cryo-EM structures. Notably, this extension can make the N-terminal region of the HTT mutant more accessible to engage in additional interactions, which is in good agreement with a previous report^38^.

Overall, these findings demonstrate that the LuTHy-BRET assay can be used to monitor dynamic changes and rearrangements (e.g. induced by mutations) within protein complexes in living cells.

## DISCUSSION

To globally understand cellular processes, it is essential to characterize complexomes to an extent where all interacting protein subunits of a complex and their relationships to one another are clearly defined. Currently, this knowledge is usually obtained when the 3D structure of a protein complex is resolved. In parallel, the Y2H system has been widely used to generate large PPI networks based on relatively simple yes/no growth readouts. However, many binary PPI assays can generate highly quantitative scores. For example, such assays have been previously used to determine relative binding affinities or to assess the effects of mutations and drugs on specific PPIs^19, 39, 40^. Here, we demonstrate that the quantitative readouts of two versatile PPI assays, LuTHy and mN2H, can be used to systematically detect protein interactions within distinct multiprotein complexes, and confidently differentiate direct PPIs from indirect associations. The presented quantitative mapping approach can thus complement high-throughput AP-MS-based and low-throughput structural biology techniques to perform systematic, in-depth characterizations of diverse protein complexes.

We provide different standards to control for the quality of such quantitative mapping efforts: 1) the published hsPRS-v2 and hsRRS-v2, which contain direct interactions and random protein pairs, respectively^18^, and 2) the direct interactions and indirect associations from three multiprotein complexes, i.e. LAMTOR, BRISC, and MIS12. These pairs of full-length proteins can systematically be introduced in future screening pipelines to calibrate PPI assays and experimentally determine scoring thresholds that will maximize detection of direct interactions, while minimizing recovery of indirect associations, inter-complex or random protein pairs. Interestingly, while LuTHy also identified indirect associations at a threshold where none of the inter-complex pairs were scored positive, the mN2H assay did not recover any of those pairs. This could potentially be explained by the fact that LuTHy assays are performed under conditions of very low donor protein expression^19^. Therefore, it is likely that endogenously expressed members of the complex might bridge the indirectly associating proteins detected by LuTHy. However, direct interactions could confidently be distinguished from indirect associations using a machine learning-based clustering algorithm that should be applicable to analyze the quantitative readouts of other binary PPI assays.

Our quantitative mapping approach should also help prioritizing high-confidence, direct interactions for further hypothesis-driven experiments such as those where PPIs are used as drug targets in chemical screens, or as candidates in mutagenesis studies. For example, identifying directly interacting subunits within disease-relevant protein complexes can pave the way for the development of small-molecule inhibitors or stabilizers of the studied complex. In addition, when a direct interaction is identified, libraries of protein fragments^41^ could easily be tested in order to identify minimal interacting domains between the two partners. This could, for example, guide structural biology efforts to solve specific 3D co-complex structures.

Since many macromolecular assemblies still remain unresolvable by traditional structural biology techniques, it is necessary to integrate complementary experimental data^42^ and theoretical approaches^43–48^. In addition, 3D structures often use truncated protein constructs and provide snapshots of the studied complexes outside their cellular environment^28, 43^. Indeed, all direct interactions occurring in live cells might not always be captured in the 3D model(s) currently available, and they are thus limited in fully representing the *in vivo* dynamics of complexes with full-length proteins. However, data obtained by binary interaction assays can provide important, complementary information on protein complex architectures in a more physiological environment. For example, single-molecule Förster resonance energy transfer (smFRET) has provided valuable results as an integrative structural biology technique by delivering information on distances between protein subunits within complexes in live cells^28, 31^. We are now adding LuTHy-BRET to the list of available integrative structural biology tools since we show that this readout can be used to measure distances between protein subunits and domains as well as to monitor dynamic conformational changes in live cells. Since LuTHy-BRET measurements are obtained from a population of molecules and not from a single molecule, it detects averaged effects within the studied cell population. Furthermore, BRET-based approaches offer three advantages over techniques like smFRET: 1) they can be applied *in vivo*^49^; 2) they have higher signal-to-noise ratios due to the absence of autofluorescence and photodestruction^50^; and 3) they do not require specialized equipment for in-cell measurements and can thus be easily implemented in any lab equipped with microtiter plate readers.

Finally, we have shown that the polyQ expansion mutation in the Huntington’s disease related HTT protein gives rise to an abnormal domain rearrangement that can be quantified in live cells with LuTHy-BRET. This suggests that this assay can potentially be used in future experiments to identify chemical modulators that reverse the effect of this and other mutations.

## AUTHOR CONTRIBUTIONS

P.T., S.G.C., J.O., M.V. and E.E.W. conceived the study. P.T., C.S., S.G.C., J.O., P.C., Y.W., E.S.R., S.G., M.Z., S.B., K.S. and Y.J. designed and performed the experiments and collected the data. P.T. analyzed the majority of the results, with significant contributions from C.S., S.G.C., J.O., M.S., T.H. while E.S.R., P.C., Y.W., J.C.T., M.A.C., D.E.H. and Y.J. provided critical insights. All authors discussed the results and P.T., C.S., S.G.C., J.O., M.V. and E.E.W. wrote the manuscript.

## ACKNOWLEDGEMENTS

The authors would like to thank all members of the Wanker, Vidal, Jacob, and Twizere laboratories for helpful discussions throughout this project. This work was supported by the Helmholtz Association, iMed and Helmholtz-Israel Initiative on Personalized Medicine (Germany); the Federal Ministry of Education and Research and e:med Systems Medicine – IntegraMent 01GS0844 (Germany) all to E.E.W.; as well as by the CHDI Foundation (USA); the German Cancer Consortium DKTK (Germany) and the Deutsche Krebshilfe, ENABLE (Germany) to E.E.W. and P.T. This work was also supported by a Claudia Adams Barr Award to S.G.C., a Fonds de la Recherche Scientifique (F.R.S.-FNRS)-Télévie Grant (FC27371, Credit no 7454518F) and a Wallonia-Brussels International (WBI)-World Excellence Fellowship to J.O. Additional supports were provided by NIH grants P50HG004233 and R01GM130885 awarded to M.V. and U41HG001715 awarded to M.V., D.E.H. and M.A.C. This work was also supported by the LabEx IBEID (grant 10-LABX-0062). M.V. is a Chercheur Qualifié Honoraire, and J.C.T. a Maître de Recherche of the Fonds de la Recherche Scientifique (F.R.S.-FNRS, Wallonia-Brussels Federation, Belgium).

## DATA AND CODE AVAILABILITY

The protein interactions from this publication have been submitted to the IMEx (http://www.imexconsortium.org) consortium through IntAct^51^ and assigned the identifier IM-29174. All UniProt and RCSB-PDB accession codes are provided in the Source Data. Python and R codes used for data analyses are available upon request.

## METHODS

### ORF sequencing and plasmid generation

For hsPRS-v2 and hsRRS-v2 proteins, the corresponding sequence-verified entry vectors published in Choi et al^18^ were Gateway cloned into the different LuTHy destination plasmids. ORFs for subunits of the LAMTOR, MIS12 and BRISC complexes were taken from the CCSB human ORFeome 8.1, which is a sequence confirmed clonal collection of human ORFs in a Gateway entry vector system^24^. In total, 16 entry plasmids were picked from the collection, single clones were isolated, and ORFs were PCR-amplified and confirmed by bi-directional Sanger DNA sequencing. Entry clones were shuttled into LuTHy and N2H destination vectors using the Gateway Cloning Technology. All resulting vectors were analyzed by PCR-amplification of cloned ORFs and DNA gel electrophoresis (N2H plasmids), or restriction digestion and sequence validation (LuTHy plasmids). For the LuTHy assay, additional control plasmids (PA-NL, Addgene #113445; PA-mCit-NL, Addgene #113444; PA-mCit, Addgene #113443; NL, Addgene #113442) were used, as previously described^16^. The pcDNA3.1 plasmids encoding HTTQ23 and HTTQ145 (glutamines encoded by CAG/CAA triplets) were gifts from the CHDI foundation that were subcloned into pDONR221 entry vectors (ThermoFisher, #12536017). The pDONR221-F8A1 entry plasmid encoding HAP40 was obtained from Source BioSciences (OCAAo5051D1091D). *HTT* and *HAP40* entry clones were shuttled into LuTHy destination plasmids, and *HAP40* was also cloned into a pDEST26-cmyc destination plasmid, which was a kind gift from Matthias Selbach. All amino acid positions in the HTT protein refer to a sequence containing 17 glutamines with a total length of 3138 amino acids. To generate the intramolecular LuTHy-BRET HTT sensors, the LuTHy plasmids pcDNA3.1 NL-HTTQ23 and pcDNA3.1 NL-HTTQ145 were digested with SfiI and SfoI that cut within the HTT sequence and the Kan/neoR resistance sequence of the plasmid backbone. The resulting larger fragment (12161 bp for Q23, and 12563 bp for Q145) were agarose gel purified using the Invisorb^®^ Fragment Cleanup kit from Invitek Molecular. Next, the HTT sequence from Q2500 to L2680 was PCR amplified using the primers 353-FWD and 341-REV. The 353-FWD primer contained a 30 bp overhang into the *HTT* sequence, and the 341-REV primer an 18 bp GSGSGS-linker sequence as well as a 7 bp overhang into the 5’-mCitrine sequence. The coding sequence for mCitrine was amplified without the start and stop codons, using the 342-FWD and the 343-REV primers. The 342-FWD and 343-REV primers both contained an 18 bp GSGSGS-linker sequence and a 7 bp overhang into the respective 5’ or 3’ *HTT* sequences. The *HTT* sequence from amino acid P2681 to the Kan/neoR cassette of the plasmid backbone was amplified using the following primers: 340-FWD and 354-REV. The 340-FWD primer contained an 18 bp GSGSGS-linker sequence and a 7 bp overhang into the 3’-mCitrine sequence, and the 354-REV primer a 30 bp overhang into the Kan/neoR cassette of the plasmid backbone. All PCR products were column purified using the MSB^®^ Spin PCRapace kit from Invitek Molecular. Finally, the digested and gel purified plasmid backbones were assembled, together with the three purified PCR products, by Gibson assembly according to the manufacturer’s protocol (New England Biolabs, E2611). The resulting colonies carrying the final plasmids were analyzed by restriction digestion and Sanger DNA sequencing. All primer sequences can be found in **Supplementary Table 8**.

### LuTHy assay procedure

The LuTHy assay was performed as previously described^19^. In brief, HEK293 cells were reversely transfected in white 96-well microtiter plates (Greiner, #655983) at a density of 4.0-4.5×10^4^ cells per well with plasmids encoding donor and acceptor proteins. After incubation for 48 h, mCitrine fluorescence was measured in intact cells (Ex/Em: 500 nm/530 nm). For LuTHy-BRET assays, coelenterazine-h (pjk, #102182) was added to a final concentration of 5 μM (5 mM stock dissolved in methanol). Next, cells were incubated for an additional 15 min and total luminescence as well as luminescences at short (370-480 nm) and long (520-570 nm) wavelengths were measured using the Infinite^®^ microplate readers M200, M1000, or M1000 PRO (Tecan). After luminescence measurements, the luminescence-based co-precipitation (LuC) assay was performed. Cells were lysed in 50-100 μL HEPES-phospho-lysis buffer (50 mM HEPES, 150 mM NaCl, 10% glycerol, 1% NP-40, 0.5% deoxycholate, 20 mM NaF, 1.5 mM MgCl_2_, 1 mM EDTA, 1 mM DTT, 1 U Benzonase, protease inhibitor cocktail (Roche, EDTA-free), 1 mM PMSF, 25 mM glycerol-2-phosphate, 1 mM sodium orthovanadate, 2 mM sodium pyrophosphate) for 30 min at 4°C. Lysates (7.5 µL) were transferred into small volume 384-well microtiter plates (Greiner, #784074) and fluorescence (mCit_IN_) was measured as previously described^19^. To measure the total luminescence (NL_IN_), 7.5 µL of 20 µM coelenterazine-h in PBS was added to each well and the plates incubated for 15 more minutes. For LuC, small volume 384-well microtiter plates (Greiner, #784074) were coated with sheep gamma globulin (Jackson ImmunoResearch, #013-000-002) in carbonate buffer (70 mM NaHCO_3_, 30 mM Na_2_CO_3_, pH 9.6) for 3 h at room temperature, and blocked with 1% BSA in carbonate buffer before being incubated overnight at 4°C with rabbit anti-sheep IgGs in carbonate buffer (Jackson ImmunoResearch, #313-005-003). 15 µL of cell lysate was incubated for 3 h at 4°C in the IgG-coated 384-well plates. Then, all wells were washed three times with lysis buffer and mCitrine fluorescence (mCit_OUT_) was measured as described^19^.

Finally, 15 µL of PBS buffer containing 10 μM coelenterazine-h was added to each well and luminescence (NL_OUT_) was measured after a 15 min incubation period.

### LuTHy data analysis

Data analysis was performed as previously described^19^. In brief, the LuTHy-BRET and LuTHy-LuC ratios from BRET and co-precipitation measurements are calculated as follows:

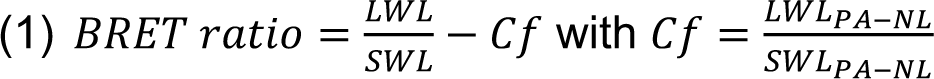

with LWL and SWL being the detected luminescences at long (520–570 nm) and short (370– 480 nm) wavelengths, respectively. The correction factor (Cf) represents the donor bleed-through value from the PA-NL only construct. The corrected BRET (cBRET) ratio is calculated by subtracting the maximum BRET ratios of control 1 (NL/PA-mCit-Y), or of control 2 (NL-X/PA-mCit) from the BRET ratio of the studied interaction (NL-X/PA-mCit-Y).

For the LuC readout, the obtained luminescence precipitation ratio (PIR) of the control protein PA-NL (PIR_PA-NL_) is used for data normalization, and is calculated as follows:

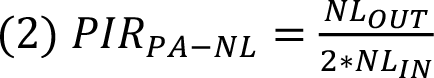

With NL_OUT_ being the total luminescence measured after co-IP and NL_IN_ the luminescence measured in the cell extracts, directly after lysis. Subsequently, LuC ratios are calculated for all interactions of interest, and normalized to the PIR_PA-NL_ ratio:

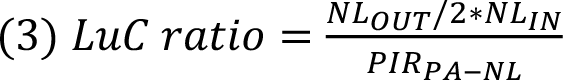

Finally, a corrected LuC (cLuC) ratio is calculated by subtracting either the LuC ratio of control 1 (NL/PA-mCit-Y), or of control 2 (NL-X/PA-mCit) from the LuC ratio of the studied interaction (NL-X/PA-mCit-Y). The calculated LuC ratios obtained for controls 1 and 2 are then compared to each other, and the highest value is used to correct the LuC ratio of the respective interaction.

### Mammalian cell-based version of the N2H assay (mN2H)

HEK293T cells were seeded at 6×10^4^ cells per well in 96-well, flat-bottom, cell culture microplates (Greiner Bio-One, #655083), and cultured in Dulbecco’s modified Eagle’s medium (DMEM) supplemented with 10% fetal calf serum at 37 °C and 5% CO_2_. 24 h later, cells were transfected with 100 ng of each N2H plasmid (pDEST-N2H-N1, -N2, -C1 or -C2) using linear polyethylenimine (PEI) to co-express proteins fused with complementary NanoLuc fragments, F1 and F2. The stock solution of PEI HCl (PEI MAX 40000; Polysciences Inc; Cat# 24765) was prepared according to the manufacturer’s instructions. Briefly, 200 mg of PEI HCl powder were added to 170 mL of water, stirred until complete dissolution, and pH was adjusted to 7 with 1 M NaOH. Water was added to obtain a final concentration of 1 mg/mL, and the stock solution was filtered through a 0.22 µm membrane. The DNA/PEI ratio used for transfection was 1:3 (mass:mass). 24 h after transfection, the culture medium was removed and 50 µL of 100x diluted NanoLuc substrate (Furimazine, Promega Nano-Glo, N1120) was added to each well of a 96-well microplate containing the transfected cells. Plates were incubated for 3 min at room temperature. Luciferase enzymatic activity was measured using a TriStar or CentroXS luminometer (Berthold; 2 s integration time).

### Processing publicly available interaction data

Publicly available binary protein interaction datasets used in this study came from the original Choi et al publication^18^. The hsPRS-v2 PPI recovery rates were extracted as published, in conditions where none of the hsRRS-v2 pairs are scored positive. To generate hsPRS-PDB, structurally supported interactions from hsPRS-v2 were identified using interactome insider (http://interactomeinsider.yulab.org), and using both, co-crystal structures and homology modeling (**Source Data Figures 1-2**). The BioPlex dataset^7,11, 14^ was downloaded from https://bioplex.hms.harvard.edu/interactions.php on May 10, 2021.

### Scoring hsPRS-v2 PPIs under conditions where no hsRRS-v2 pair is scored positive for LuTHy

Source data for each assay version performed in this study are presented in the **Source Data Figures 1-2**. Each binary PPI experiment with LuTHy-BRET and LuTHy-LuC was performed six times, with two biological and three technical replicates. For each protein pair X-Y, we calculated the corrected BRET and LuC ratios as described above. To identify protein pairs that scored positive at a threshold of no hsRRS-v2 detection for a given tagging configuration, only cBRET or cLuC ratios higher than the highest hsRRS-v2 score for that tagging configuration were considered. All hsPRS-v2 pairs that did not meet this criterion were not scored positive and therefore defined as not detected in the corresponding assay version.

### Complex selection and definition of direct interactions and indirect associations

Human protein complexes used in this study were selected based on the following criteria. First, human protein complexes should have at least one experimentally determined structure in PDB^23^ (https://www.rcsb.org). Second, the complex should have at least four subunits. Third, at least 80% of entry clones for individual subunits of a complex should be present in the human ORFeome 8.1 collection^24^. A total of 24 distinct complexes (**Supplementary Table 1**) with different PDB structures met those criteria, and three protein complexes with well-documented biological functions were selected from this list: LAMTOR, BRISC and MIS12. These complexes were also picked as they appear in AP-MS interactome maps, according to BioPlex^7,11, 14^. Finally, direct interactions and indirect associations within protein complexes were determined by interactome 3D^25^ and PDBePISA^26^.

### Scoring interactions within multiprotein complexes

Data for LuTHy and mN2H mapping of multiprotein complexes can be found in **Source Data Figures 3-5**. Recovery of interactions by LuTHy-BRET, LuTHy-LuC and mN2H was calculated based on construct-specific cutoffs. Therefore, the quantitative scores were determined as described in the LuTHy and mN2H methods sections, respectively. For each construct, X or Y, involved in the X-Y interaction, a cutoff was defined as the highest inter-complex pair score for each of the two respective constructs (X and Y). The final cutoff for a tested protein pair between constructs X and Y was then determined as the maximum of the two cutoff values.

### Cluster analysis of LuTHy data

For supervised classification of direct and not-direct interaction clusters, a Gaussian finite mixture model using the R package mclust^53^ was used. First, the LuTHy-BRET and LuTHy-LuC quantitative scores (i.e. cBRET and cLuC ratios) from 63 LuTHy-positive interactions among hsPRS-PDB and from 585 protein pairs in hsRRS-v2 (**Supplementary Table 3**) were used as positive and negative training sets, respectively. The MclustDA discriminant analysis function was applied to those training sets using the eigenvalue decomposition discriminant analysis (EDDA) method. The training set performed at a 2.36% classification error rate. Next, the trained algorithm was used to cluster 117 intra-complex protein pairs (**Supplementary Table 3**) that scored above the construct-specific cutoffs in LuTHy assays. Using the function predict.MclustDA, 47 direct protein pairs (i.e. multiple configurations corresponding to 14 out of the 23 LuTHy-positive direct interactions) were correctly classified as direct interactions and all 12 indirect protein pairs (i.e. multiple configuration corresponding to all 6 LuTHy-positive indirect associations) were accurately classified as not-direct. In addition, 58 direct protein pairs (i.e. multiple configuration corresponding to 9 out of the 23 LuTHy-positive direct interactions) could not be classified as direct interactions using this supervised classification analysis.

### Determining interaction interface areas, complex surface areas and molecular distances between protein termini

Interaction interface and total complex surface areas were determined according to PDBePISA^26^ (https://www.ebi.ac.uk/msd-srv/prot_int/pistart.html) on April 16, 2021. Protean 3DTM (DNASTAR^®^) was used to measure molecular distances between protein termini of interactions in the hsPRS-PDB as well as between direct interactions and indirect associations within the multiprotein complexes: LAMTOR (RCSB PDB: 6EHR), MIS12 (RCSB PDB: 5LSJ) and BRISC (RCSB PDB: 6H3C). The determined molecular distances can be found in **Source Data Figures 3-5**. Nonlinear and linear regressions between cBRET ratios and molecular distances between protein termini were calculated using GraphPad Prism 9 and the fits were compared using the extra-sum-of-squares F test with a bottom constraint to 0. GraphPad Prism 9 was also used to interpolate molecular distances and their 90% confidence intervals using the measured cBRET ratios.

---

**Supplementary Figure 1 (related to Figure 1).**
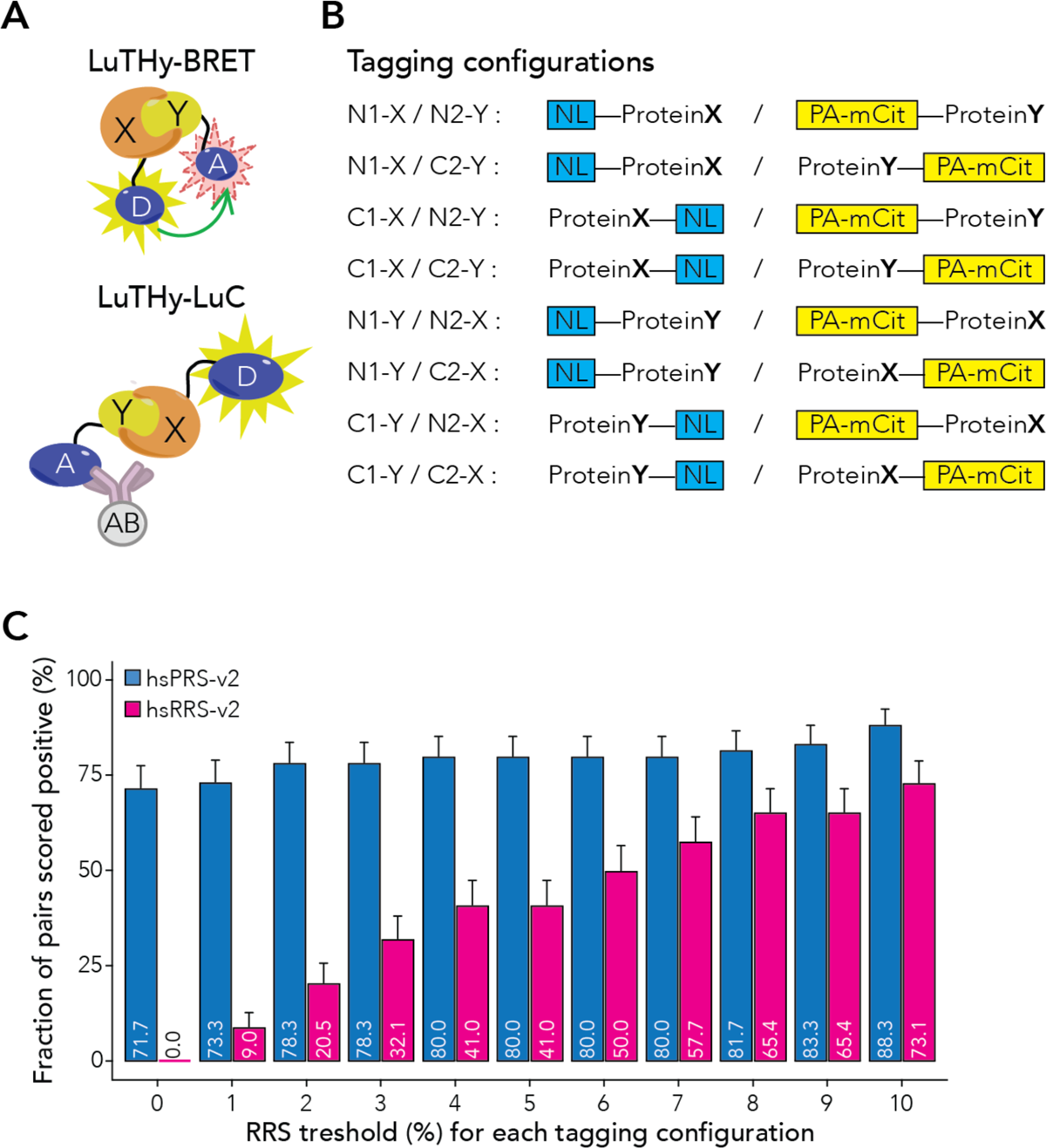
Tagging configurations and relationship between the hsRRS-v2 threshold and detection of hsPRS-v2 and hsRRS-v2 pairs for LuTHy. (**A**) Schematic overview of the LuTHy-BRET and LuTHy-LuC assays. X: Protein X, Y: Protein Y, D: NanoLuc donor, A: mCitrine acceptor, AB: antibody. (**B**) With the LuTHy assay, each protein pair X-Y can be tested in four possible configurations (N- vs. C-terminal fusion for each protein), and proteins can be swapped from one tag to the other for a total of 8 possible LuTHy versions resulting in 16 quantitative scores for each protein pair (i.e. eight for LuTHy-BRET and eight for LuTHy-LuC). (**C**) Impact of scoring random protein pairs on PPI recovery for all LuTHy versions: cumulative detection rates of hsPRS-v2 and hsRRS-v2 pairs when increasing, identical hsRRS-v2 thresholds are applied for each individual assay version. Error bars in (**C**) indicate standard errors of the proportion.

**Supplementary Figure 2 (related to Figure 1).**
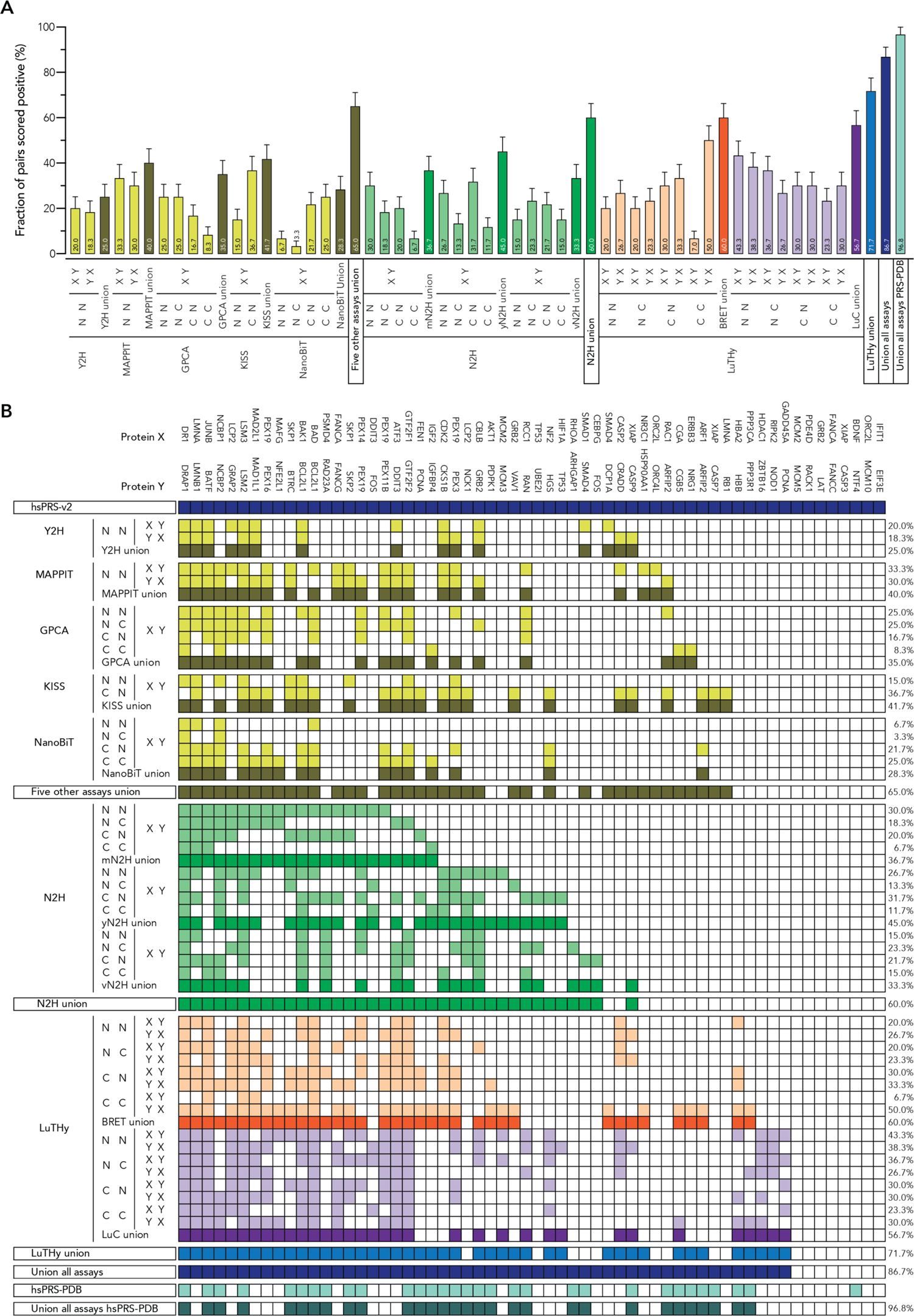
Performance and complementarity of different binary PPI assay versions under conditions where none of the hsRRS-v2 pairs are scored positive. (**A,B**) Performance of individual binary PPI assay versions benchmarked in Choi et al^18^ (Y2H, MAPPIT, GPCA, KISS, NanoBiT, N2H), and in this study (LuTHy). In total, 42 assay versions were tested, reaching ∼87% detection of hsPRS-v2 PPIs. The 31 hsPRS-PDB pairs correspond to hsPRS-v2 PPIs currently supported by at least one 3D structure in PDB. The percentage (%) at the end of each row represents the fraction of hsPRS-v2 PPIs scored positive when none of the hsRRS-v2 pairs are recovered. Error bars in (**A**) indicate standard errors of the proportion.

**Supplementary Figure 3 (related to Figure 3).**
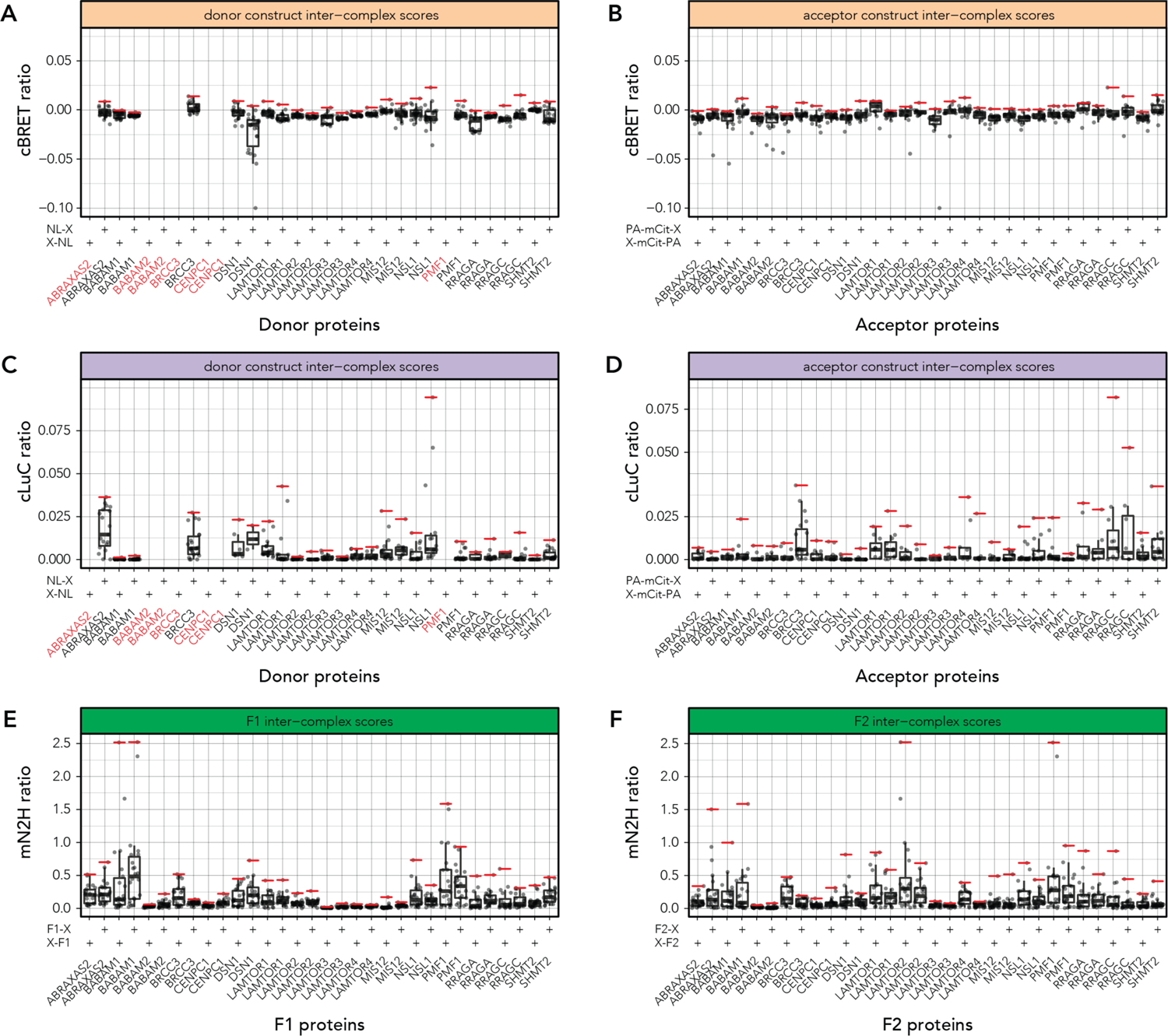
Establishing a construct-specific scoring threshold. For each assay and expression construct, boxes and whiskers for the quantitative scores of inter-complex pairs are shown (displayed are the median, lower and upper hinges showing 25^th^ and 75^th^ percentiles, lower and upper whiskers extending from the hinges with 1.5x the inter-quartile range). Individual scores are indicated by small grey dots. The horizontal red lines indicate the cutoffs applied for each construct. Constructs written in red on the x-axes showed no detectable expression. Scores of inter-complex pairs differentiated by individual constructs for LuTHy-BRET (**A,B**), LuTHy-LuC (**C,D**), and mN2H (**E,F**) assays. Each protein pair was tested in every possible configuration (i.e. C-C, C-N, N-C, N-N), and each protein was also tested as donor (**A, C**) and acceptor (**B, D**) in LuTHy, or as F1 (**E**) and F2 (**F**) NanoLuc fusions in mN2H. The (+) indicates the N- or C-terminal tagging configuration of the indicated protein.

**Supplementary Figure 4 (related to Figure 3).**
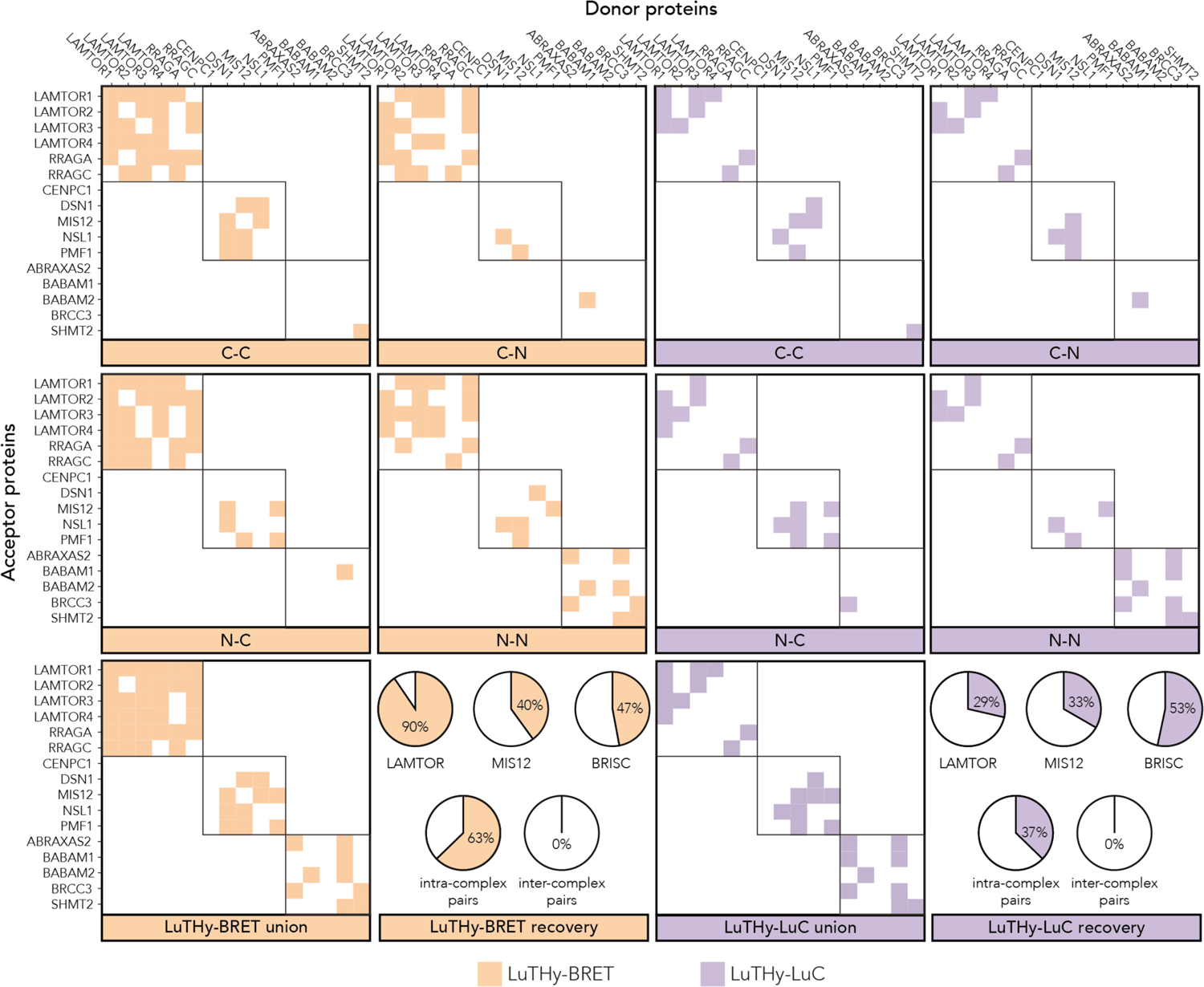
Mapping interactions within multiprotein complexes using the LuTHy-BRET and LuTHy-LuC assays. Results of the all-by-all interaction screen with the LuTHy-BRET and LuTHy-LuC assays for the selected multiprotein complexes. Each protein pair was tested in every possible configuration (i.e. C-C, C-N, N-C, N-N), and each protein was also tested as donor and acceptor. In the pie charts, the top panels show the recovery rates of intra-complex pairs within the LAMTOR, MIS12 and BRISC complexes, while the bottom panels show the recovery rates of all intra-complex and inter-complex pairs.

**Supplementary Figure 5 (related to Figure 3 and 4).**
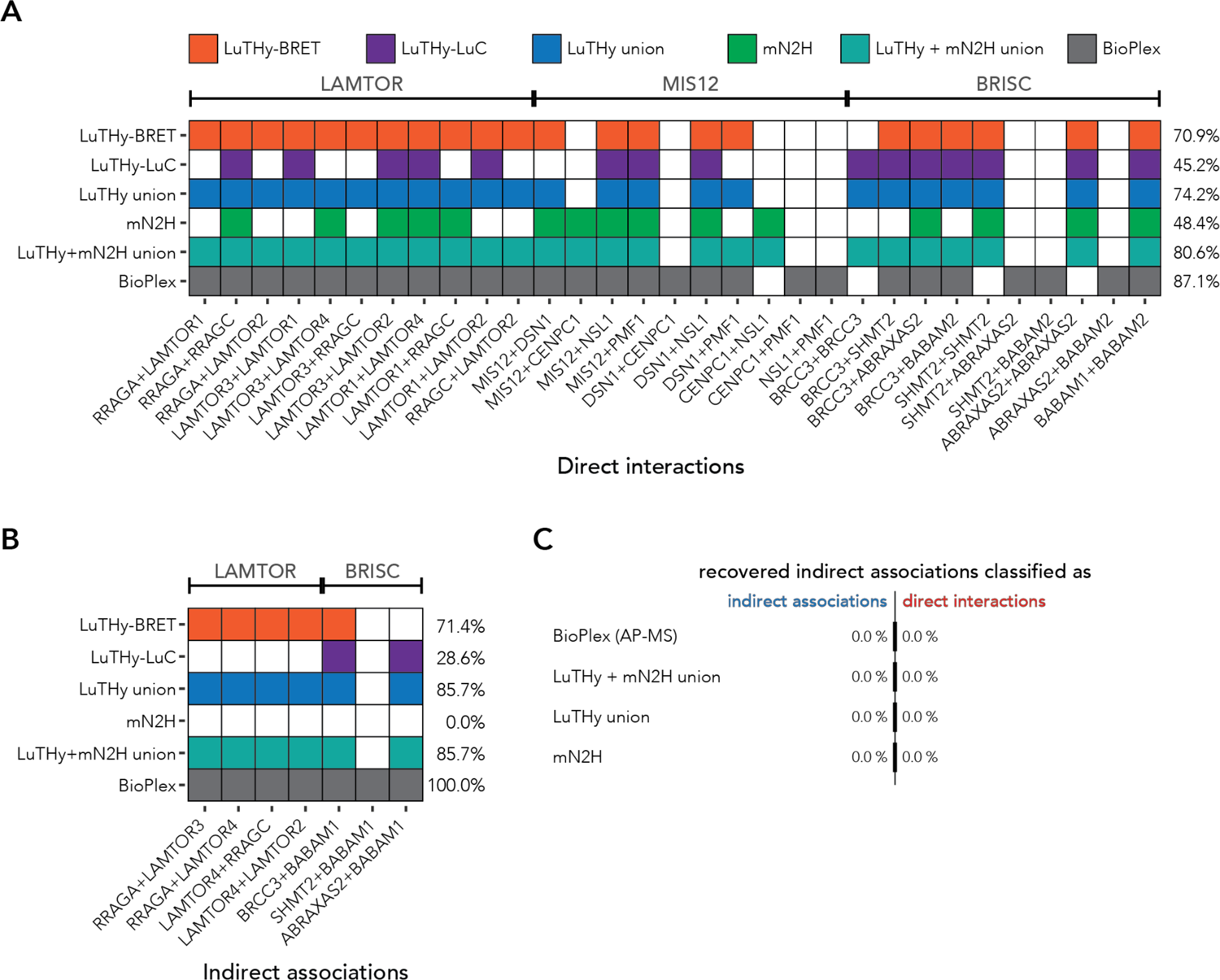
Summary of the direct interactions and indirect associations detected by different approaches. Detected direct interactions (**A**) and indirect associations (**B**) by the LuTHy and mN2H binary PPI assays, and by AP-MS-based (BioPlex) techniques. The percentage (%) at the end of each row represents the fraction of protein pairs recovered (for LuTHy and mN2H: at a threshold where no inter-complex pair is scored positive). LuTHy + mN2H union summarizes the LuTHy and mN2H results. Published reference data from the AP-MS BioPlex dataset are shown. Structural biology data were used to define direct interactions and indirect associations. (**C**) Recovery and classification of structurally defined indirect associations as true indirect associations or as direct interactions for BioPlex, LuTHy and mN2H.

**Supplementary Figure 6 (related to Figure 4).**
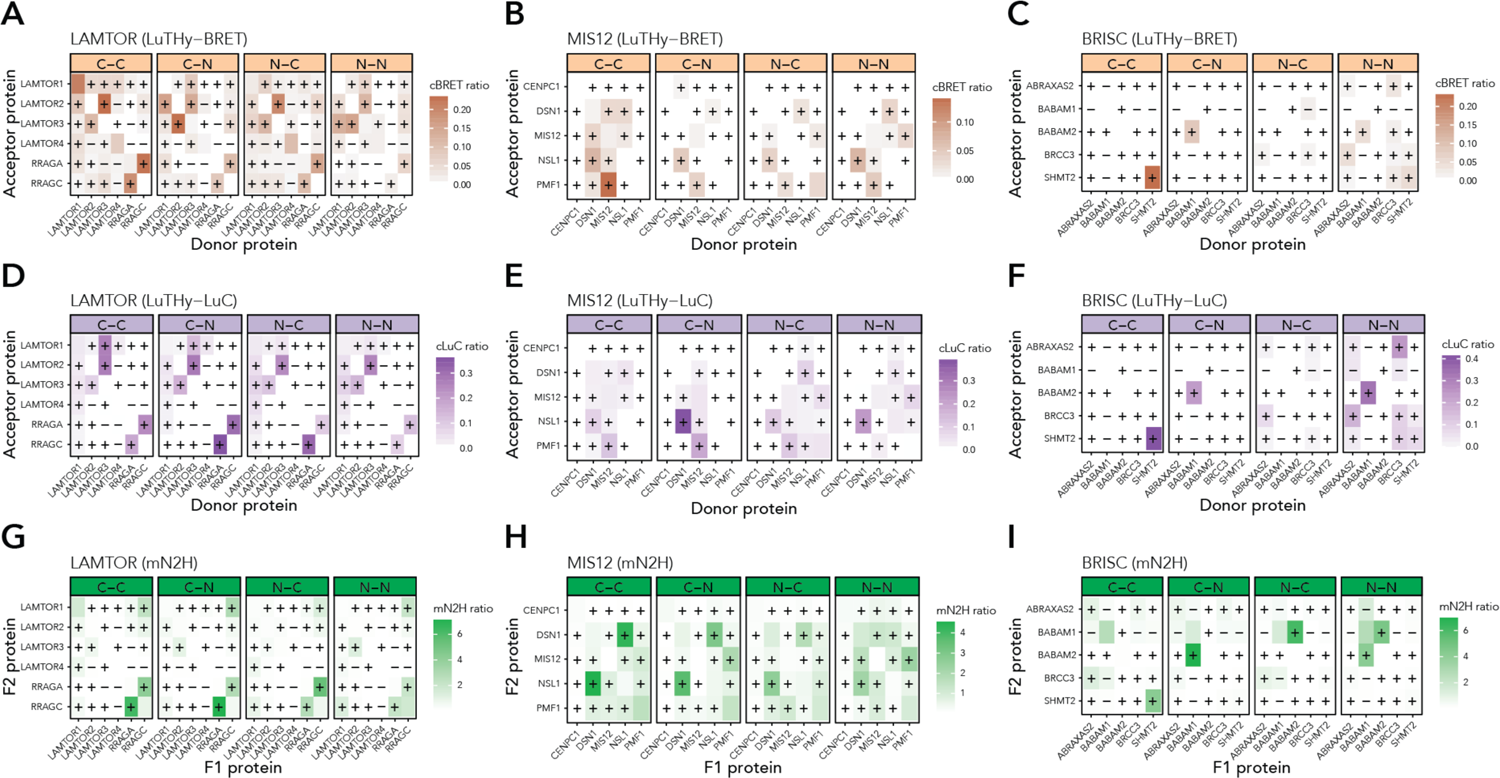
Quantitative scores for direct interactions and indirect associations according to the tested tagging configurations. Heat maps of all intra-complex protein pairs in the LAMTOR, MIS12 and BRISC complexes, differentiated by tagging configurations for LuTHy-BRET (**A-C**), LuTHy-LuC (**D-F**) and mN2H (**G-I**). The range of quantitative scores is depicted from smaller, or equal to zero (white) to maximal (orange, purple or green for LuTHy-BRET, LuTHy-LuC or mN2H, respectively) values. A “+” symbol indicates a direct interaction, a “-” symbol an indirect association, and an empty cell a separate homodimer interaction.

**Supplementary Figure 7 (related to Figure 4).**
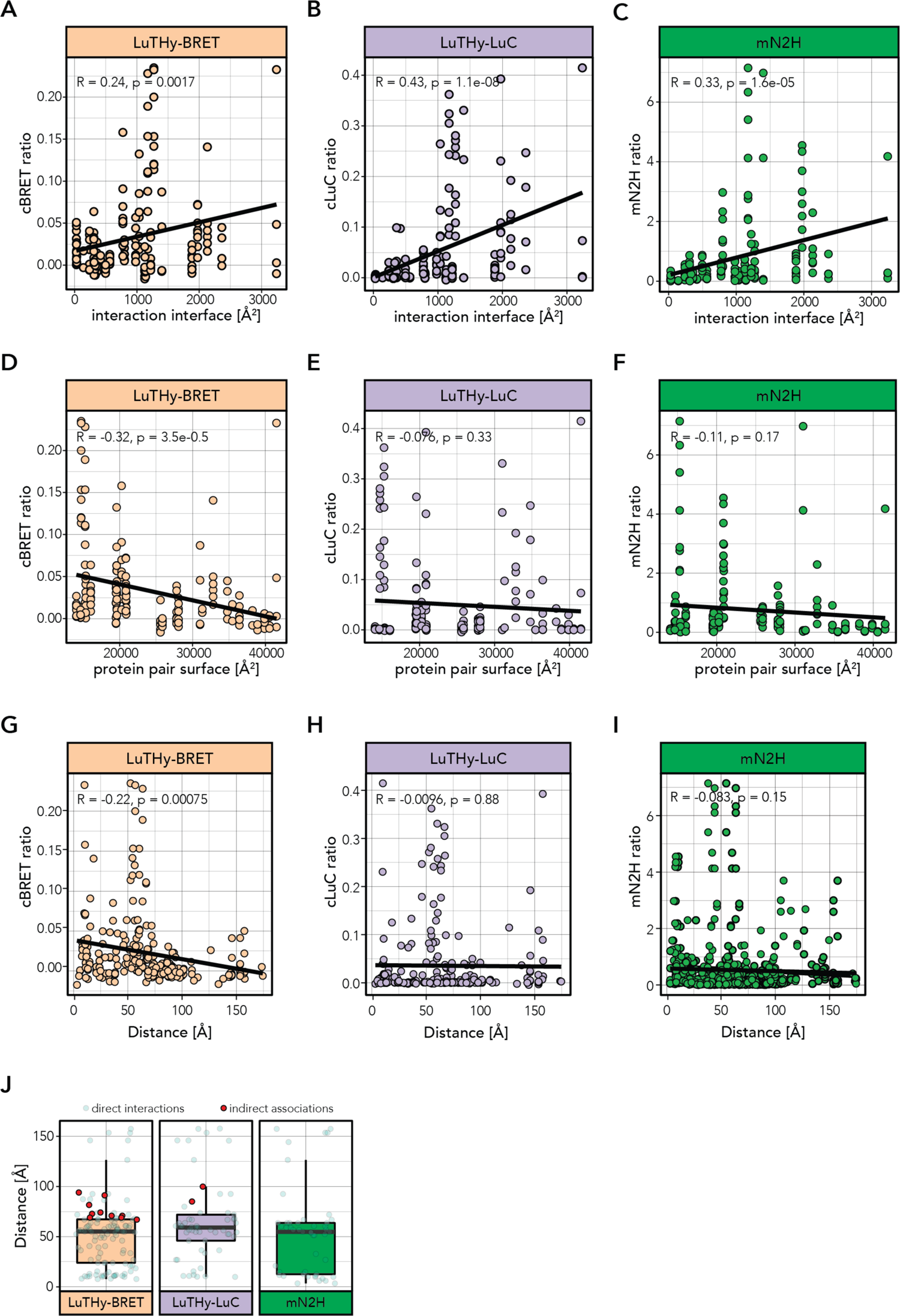
Correlation between the LuTHy and mN2H quantitative scores and the structural properties of the analyzed complexes. Correlation between the interaction interfaces between proteins and the cBRET (**A**), cLuC (**B**), and mN2H (**C**) ratios for the corresponding PPIs. Correlation between the total surface areas of interacting proteins and the cBRET (**D**), cLuC (**E**), and mN2H (**F**) ratios. Correlation between the molecular distances (Å) of the tagged protein termini and the cBRET (**G**), cLuC (**H**), and mN2H (**I**) ratios of the corresponding PPIs. Pearson correlation coefficients (R) with p-values are displayed. (**J**) Box plots of the molecular distances (Å) between the tagged protein termini for the detected direct interactions (transparent green) and indirect associations (red) in the LuTHy-BRET, LuTHy-LuC and mN2H assays (displayed are the median, lower and upper hinges showing the 25^th^ and 75^th^ percentiles, lower and upper whiskers extending from the hinges with 1.5x the inter-quartile range).

**Supplementary Figure 8 (related to Figure 4).**
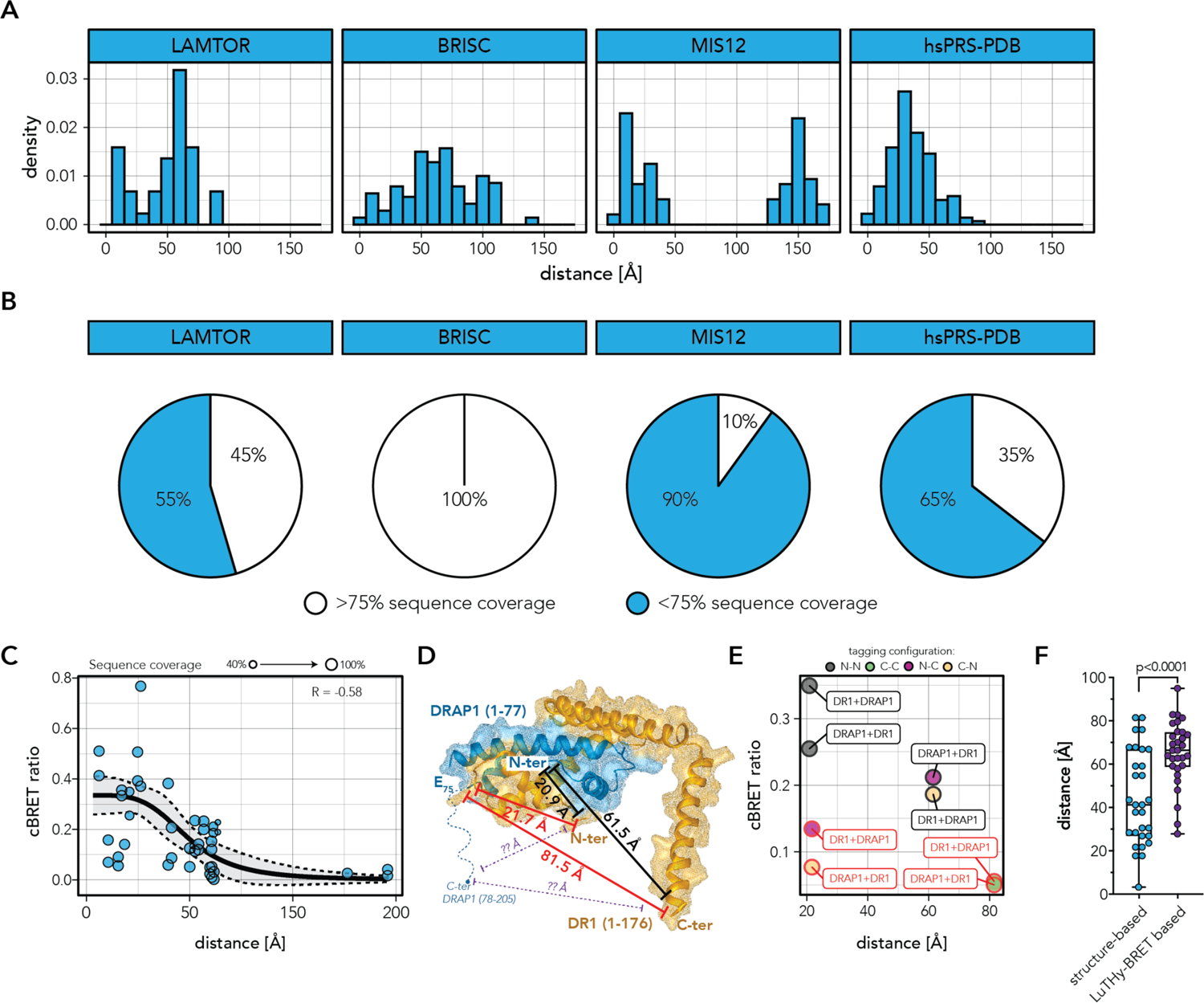
Protein sequence coverage and molecular distances between subunits of the studied complexes used to generate the cBRET-distance standard curve. (**A**) Relative distribution of structure-based distances for direct interactions within the LAMTOR, BRISC and MIS12 complexes, as well as in the hsPRS-PDB PPIs. (**B**) Proportion of directly interacting proteins reported in the 3D structures that contain, on average, more (white) or less (green) than 75% of the full-length protein sequences. (**C**) Correlation between the cBRET ratios and the molecular distances for the 44 protein pairs where at least six tagging configurations out of the eight tested are scored positive in LuTHy-BRET, and for which the respective tagged protein termini are structurally resolved. Goodness of the sigmoidal regression fit is indicated on the graph (R = −0.58). (**D**) 3D structure of the DR1-DRAP1 interaction where molecular distances between the last amino acids of the structurally solved proteins are indicated in red. The dotted blue lines indicate the protein fractions that are not structurally resolved (DRAP1 C-terminus). Unknown molecular distances between the missing termini in the 3D structure are indicated by dotted purple lines. (**E**) cBRET ratios for the DR1-DRAP1 interaction are plotted against the structure-based molecular distances. Tagging configurations are colored by N-N (grey), C-C (green), N-C (purple) or C-N (yellow). Each data point on the graph is labeled (framed text) according to the tested tagging configuration: the protein indicated first is tagged with NanoLuc (NL) luciferase, while the second protein is tagged with PA-mCitrine (PA-mCit) (e.g. DR1+DRAP1 (N-N, grey) corresponds to NL-DR1/PA-mCit-DRAP1). Tagging configurations where tags are fused to the structurally unresolved termini in the current 3D structure for one of the two proteins are outlined in red (e.g. DR1+DRAP1 (N-C, purple) corresponds to NL-DR1/DRAP1-mCit-PA). (**F**) Structure-based and predicted distances for the 30 PPIs with tags fused to protein termini not currently resolved in the structures (box and whisker visualizing the median, lower and upper hinges showing the 25^th^ and 75^th^ percentiles, lower and upper whiskers extending from min to max; all data points are shown). Statistical significance was calculated by a two-tailed, paired t-test, n=30.

**Supplementary Figure 9 (related to Figure 4).**
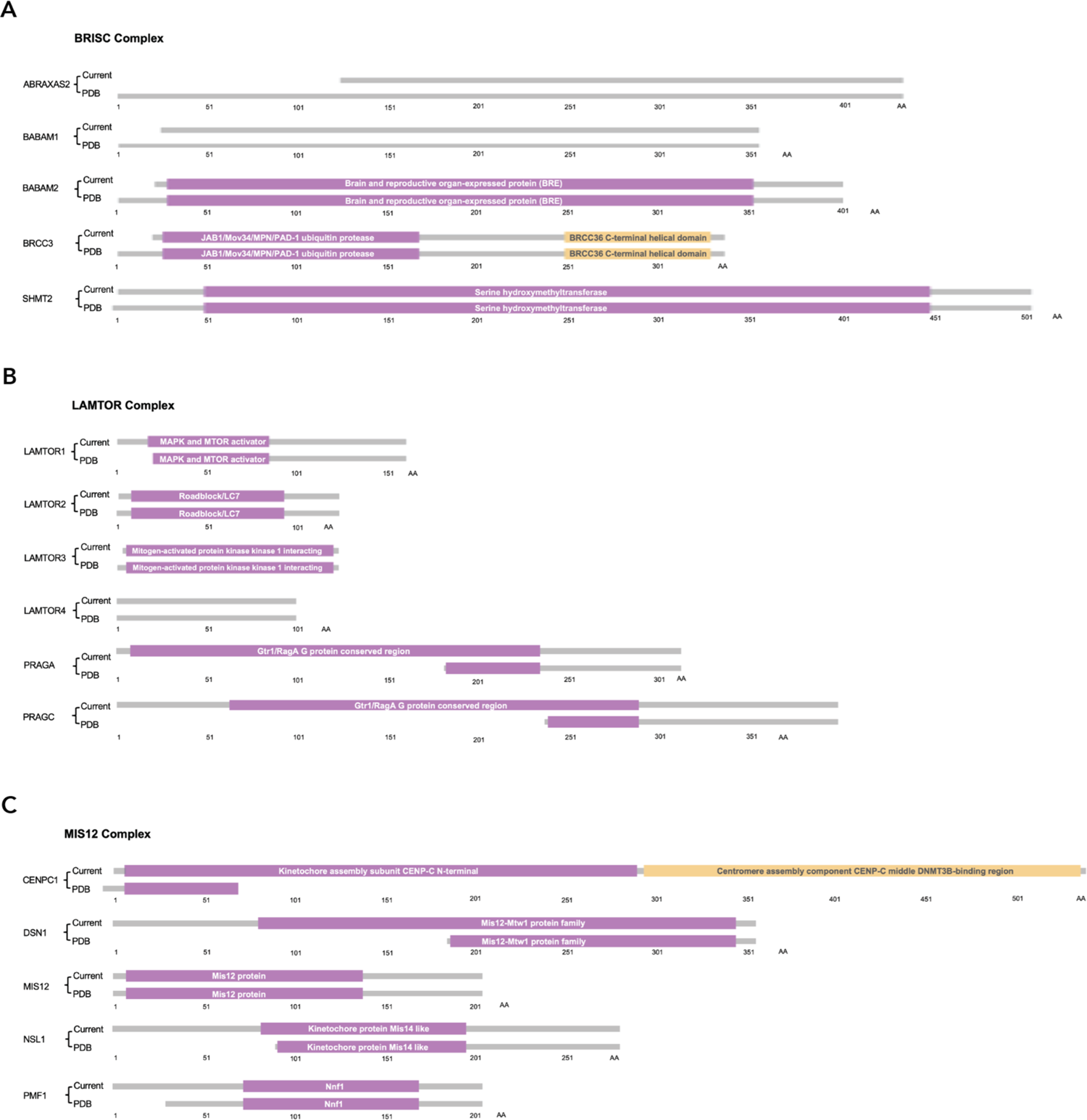
Protein sequence coverage of the constructs used in the present binary interaction study (“Current”) compared to those found in the published structures (“PDB”). Schematic representations of the protein constructs used in the BRISC (**A**), LAMTOR (**B**), and MIS12 (**C**) complexes. The numbers indicate the lengths of the protein constructs in amino acids (AA). (**A**) The structural study characterizing the BRISC complex (PDB: 6H3C) used full-length proteins whereas four of the constructs (ABRAXAS2, BABAM1, BABAM2, BRCC3) used in the current study are C-terminally truncated. The current binary interaction study uses proteins containing >90% of the BABAM1, BABAM2 and BRCC3, and ∼80% of the ABRAXAS2 full-length reference sequences. (**B**) The structural study characterizing the LAMTOR complex (PDB: 6EHR) uses smaller fragments for the RRAGA and RRAGC subunits. Except for a single AA missing on the N-terminus of LAMTOR3, the current study uses the full-length sequences for subunits of the LAMTOR complex. (**C**) Overall, the structural study of the MIS12 complex (PDB: 5LSJ) lacks >50% of the AA sequences for the different subunits.

**Supplementary Figure 10 (related to Figure 4).**
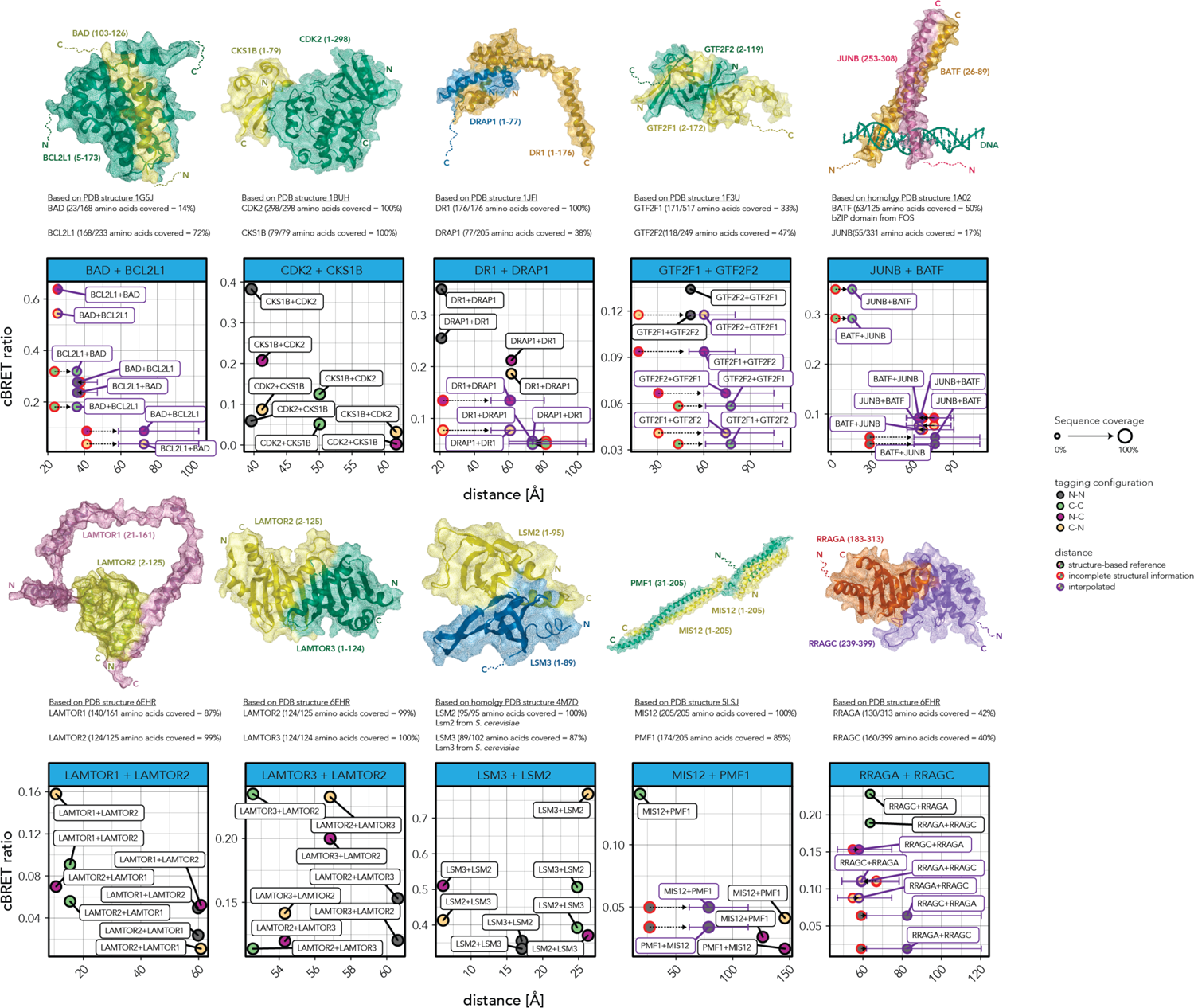
Subset of interactions where at least six out of the eight tested tagging configurations scored positive in the LuTHy-BRET assay. The following 3D structures are represented: 1G5J (BAD-BCL2L1), 1BUH (CDK2-CKS1B), 1JFI (DR1-DRAP1), 1F3U (GTF2F1-GTF2F2), 1A02 (JUNB-BATF), 6EHR (LAMTOR1-LAMTOR2 and LAMTOR3-LAMTOR2), 4M7D (LSM3-LSM2), 5LSJ (MIS12-PMF1), and 6EHR (RRAGA-RRAGC). On each structure, the protein termini are indicated by N or C. Protein sequence coverages used in the structures are also indicated (%). Protein regions that are missing in the structures are indicated by dotted lines. For all interactions, cBRET ratios are plotted against the molecular distances measured in the corresponding 3D structures or interpolated using the cBRET standard curve. The quantitative scores outlined in black correspond to tagging configurations where the tagged protein termini are resolved in the structures. These scores were used as reference data points for distance interpolations. Interaction scores and structure-based molecular distances for tagging configurations where the tagged protein termini are absent from the 3D structure are outlined in red. Interaction scores outlined in purple correspond to cBRET ratios against interpolated molecular distances. Changes in structurally measured distances versus the interpolated distances are indicated by dotted arrows. The purple horizontal error bars correspond to the 95% confidence intervals of the interpolated molecular distances. Tagging configurations are color-coded: N-N (grey), C-C (green), N-C (purple), and C-N (yellow). Each data point on the graph is labeled (blank frame) according to the associated tagging configuration tested: the protein indicated first is tagged with NanoLuc (NL) luciferase, while the second protein is tagged with PA-mCitrine (PA-mCit) (e.g. DR1-DRAP1 (N-N, grey) corresponds to NL-DR1/PA-mCit-DRAP1). Tagging configurations where tags are fused to the missing termini in the current 3D structure for one of the two proteins are outlined in red (e.g. DR1-DRAP1 (N-C, purple) corresponds to NL-DR1/DRAP1-mCit-PA). The size of the dots indicates the average percentage of protein sequence coverage in the corresponding 3D structure of the PPI.

## APPENDIX

Supplementary Table 1: List of the multiprotein complexes meeting the prioritization criteria in this study. Supplementary Table 2: Number of inter-complex pairs, direct interactions and indirect associations for the three selected protein complexes: LAMTOR, BRISC and MIS12 as defined by their 3D structures. The numbers of separate homodimer interactions not reported in the structures are also indicated.

Supplementary Table 3:

Sheet 1 - training sets: Positive and negative training sets for LuTHy cluster analysis.

Sheet 2 - clustering results: Supervised classification of direct and not-direct interaction clusters.

Supplementary Table 4: Surface and interface areas (Å2) between directly interacting proteins in the LAMTOR, BRISC and MIS12 complexes.

Supplementary Table 5: Names of protein complexes, protein subunits, and PDB IDs used in this study. The amino acid sequences of the proteins used for binary PPI mapping or in the PDB structures are indicated.

Supplementary Table 6. Characterization of protein family (pfam) domains.

Supplementary Table 7: Structure-based and BRET-interpolated distances for the HTT-HAP40 interaction, and for the intramolecular HTT sensors for HTTQ23 and HTTQ145.

Supplementary Table 8: Primer sequences used in this study.

Supplementary Table 9: Statistical reports for relevant figures.

Source Data Figures 1-2: Raw LuTHy data for interactions in hsPRS-v2.

Source Data Figures 3-5:

Sheet 1 - Multi-protein complexes: Raw LuTHy and mN2H mapping data for the selected multiprotein complexes. Molecular distances measured between structurally supported interactions between protein pairs within the LAMTOR, BRISC and MIS12 complexes.

Sheet 2 - hsPRS-PDB: Molecular distances measured between structurally supported heterodimer interactions of the hsPRS-v2 (hsPRS-PDB).

